# Whole-genome single-cell multimodal history tracing to reveal cell identity transition

**DOI:** 10.1101/2025.08.12.669973

**Authors:** Yumiko K. Kawamura, Valentina Khalil, Taro Kitazawa

**Author notes:** These authors contributed equally to this work (Yumiko K. Kawamura, Valentina Khalil).

## Abstract

Advances in single-cell sequencing have deepened our understanding of cellular identities. However, because they inherently capture only static snapshots, after which no further observations are possible, we cannot compare past and present profiles within the same cell. Thus, multi-time-point whole-genome profiling at single-cell resolution has been a long-standing goal. Here, we introduce the History Tracing-sequencing (HisTrac-seq) platform, which enzymatically labels genomic DNA adenine to “bookmark” gene regulatory statuses. This first enabled the profiling of transcriptomic and epigenetic states in the mouse brain over a period of two months. Furthermore, extending HisTrac-seq to single-cell multi-omics sequencing, we demonstrated the simultaneous mapping of past and present profiles of the same single cells. Analyzing over 93,000 cells, we discovered unexpected, drastic cell identity transitions on a large scale (“identity jumps”). This phenomenon was previously unobservable with current technologies and revealed a hidden layer of developmental plasticity. HisTrac-seq offers a powerful approach to “temporal multi-omics” for disentangling dynamic biological processes involved in development, plasticity, aging, and disease progression.

## INTRODUCTION

Recent advances in single-cell (sc) sequencing, including transcriptomics and epigenomics, have dramatically expanded our view of molecular heterogeneity and cell identity regulation in complex tissues, which are vital for appropriate development and environmental adaptation^1–6^. However, because these methods require cell lysis to harvest nucleic acids, they capture only static snapshots at each time point. Thus, even when sampling multiple time points, it remains challenging to unambiguously trace the molecular trajectories of individual cells. Trajectory-inference algorithms depend on continuity between single-cell snapshots, making them ill-suited to reconstruct processes that are highly disruptive or span long timescales^7,8^.

Although several history-tracing methods have been proposed^9–30^, multi–time-point single-cell profiling of whole-genome signatures remains highly challenging. CRISPR-based barcoding systems (e.g., DNA Typewriter^11^ coupled with ENGRAM (enhancer-driven genomic recording of transcriptional activity in multiplex)^12^ can sequentially inscribe enhancer-activation events onto a synthetic “DNA Tape.” However, it is restricted to bulk readouts, tracks only fewer than one hundred pre-selected enhancer elements, and remains untested *in vivo.* An enzymatic DNA-modification approach, DNA cytosine methyltransferase time machine (DCM-TM), fuses a DCM to RNA polymerase II (RNAPII) to deposit unique cytosine methylation marks at transcribed loci for retrospective, genome-wide mapping of gene activity *in vivo*^14^. However, DCM-TM also depends on bulk tissue analysis and cannot capture other essential chromatin modalities. Live-seq, by contrast, enables repeated sampling of cytoplasmic RNA from the same single cell *in vitro*^13^. But it requires continuous optical and physical access, offers low throughput (i.e., up to several hundred cells), remains hardly applicable *in vivo*, and cannot harvest nuclei for parallel epigenetic analyses. Consequently, an approach that combines genuine temporal resolution with genome-wide, single-cell multi-omics depth has been sought.

Here, we present HisTrac-seq, a whole-genome, multimodal history-tracing platform based on DNA adenine methyltransferase identification (DamID)^31–35^. By transiently activating DNA adenine methyltransferase (Dam) fused to a protein of interest (POI), we bookmark POI-interacting GATC motifs with N⁶-methyladenine (m6A), a modification absent in mammalian genomic DNA^31–35^. This genomic DNA labelling enables retrospective mapping of past molecular profiles in postmitotic neurons and cardiomyocytes. Swapping POIs allows interrogation of multimodal regulatory layers, including transcriptome, chromatin accessibility, epigenetic suppression, and bivalent poised domains. We demonstrate that HisTrac-seq is applicable over two months *in vivo*. Furthermore, we developed scHisTrac-seq based on the 10x Genomics Chromium platform. This simultaneously maps past and present multi-omics profiles from the same single-cells. By analysing the development of 93,844 single-cells across various epigenetic and transcriptional modalities, we uncovered large-scale, drastic cell identity transitions, namely “cell identity jumps,” phenomena that elude conventional snapshot approaches. We also delineated the epigenetic and transcriptional underpinnings of identity jumps, which are linked to premature activation of developmental programs. This sheds a new insight into the potential hidden cell identity plasticity during development, opening new avenues for dissecting how abrupt identity transitions shape tissue formation and function. Together, these findings underscore HisTrac-seq’s approach for dissecting dynamic biological processes involved in development, memory formation, aging, and disease progression.

## RESULTS

### Epigenetic and transcriptional multimodal labelling system

Previous studies of DamID have shown that m6A labelling can persist on DNA for several hours, lasting until the next round of DNA replication, although these investigations were limited to repressive signatures (e.g., Polycomb, heterochromatin) in *in vitro* culture^35,36^. Due to the absence of active adenine demethylation machinery, we hypothesized that m6A potentially serves as a robust “bookmark” for long-term recording to capture a broad spectrum of gene regulatory mechanisms in postmitotic cells (Fig. 1a). We generated seven Dam-POI fusions, five previously validated for DamID (RNAPII, Free-Dam, Laminb1, Taf3-Phd domain, and anti-H3K27me3 mintbody)^37,38^, and two novel constructs (Leo1 transcription elongation factor and a fusion of Taf3-Phd domain and Cdh7-chromodomain to label bivalency^39^) (Fig. 1b).

**Fig 1.**
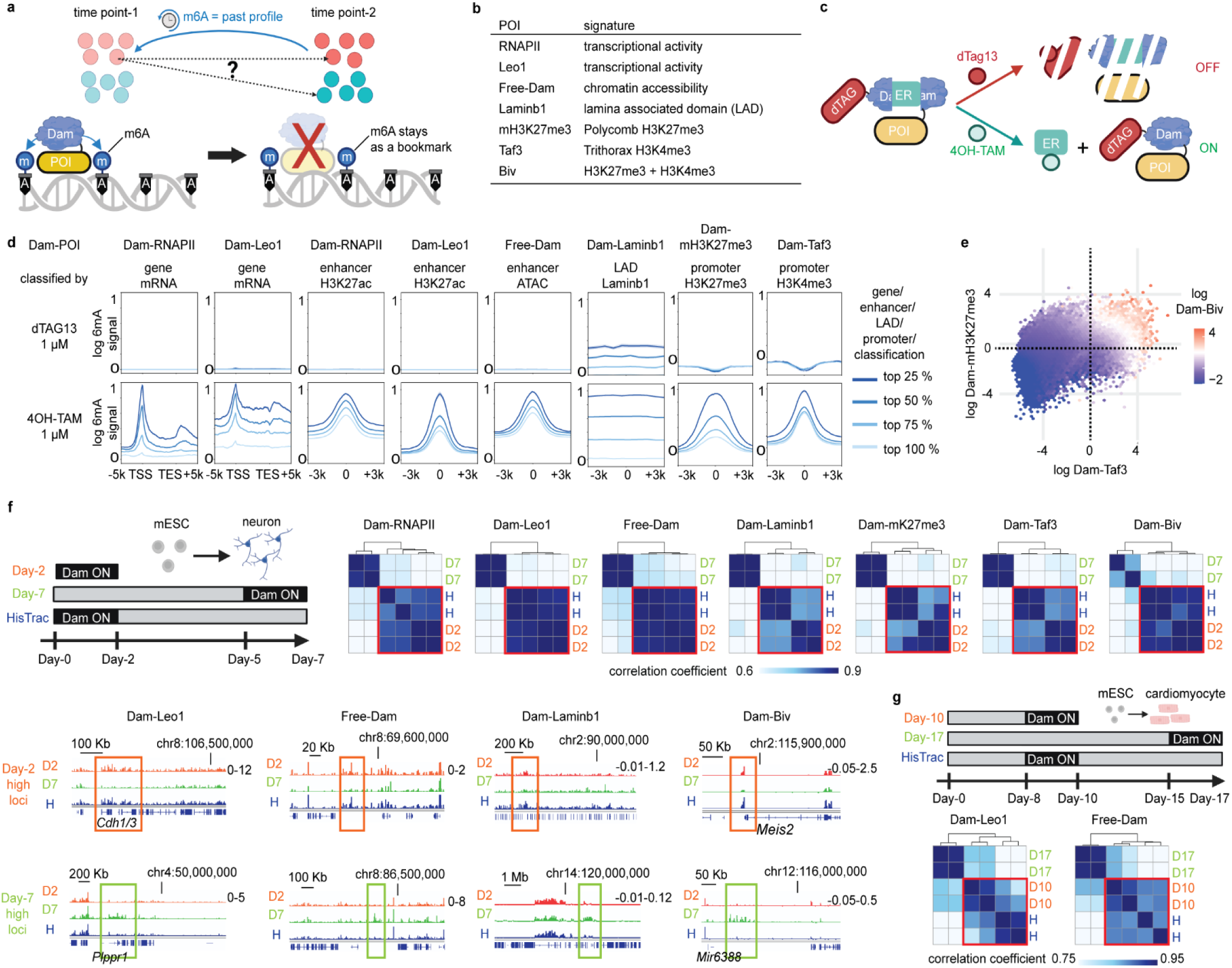
HisTrac-seq for whole-genome multimodal history tracing. **a**, Overview of HisTrac-seq. Transient activation of Dam fused to a protein of interest (POI) deposits m6A “bookmarks” at POI-bound genomic loci, enabling retrospective mapping of past molecular states of each cell. **b**, List of seven Dam-POI constructs and their targets. **c**, Temporal control of Dam activity. Dam is activated by 4OH-TAM and inactivated upon dTAG-13 treatment. **d**, Meta-analysis of m6A distribution in mESCs under Dam-off (dTAG-13+) and Dam-on (4OH-TAM+) conditions (*n* = 2). Genomic features (genes, enhancers, LADs, promoters) were stratified into quartiles (25%, 50%, 75%, 100%) based on their respective activity marks (mRNA-seq, H3K27ac-ChIP, ATAC-seq, Laminb1-ChIP, H3K27me3-ChIP, H3K4me3-ChIP). Curves show mean m6A signal ± s.e.m. **e**, Hexbin plot of Dam-Taf3 versus Dam-mH3K27me3 signal in mESCs, colored by Dam-Biv enrichment. Quantified in 10 Kb genomic bins (*n* = 2). **f**, *In vitro* neurodifferentiation history tracing (n = 2). (top left) Timeline of mESC differentiation to neurons with Dam-POI activation windows for Day-2, Day-7, and retrospective HisTrac (Day-2 -> Day-7) labelling. (top right) Genome-wide m6A quantification in fixed genomic bins clustered by Pearson’s correlation. (bottom) Genome browser tracks of Day-2- and Day-7-enriched loci for Dam-Leo1, Free-Dam, Dam-Lamin B1, and Dam-Biv, highlighting stage-specific bookmarking. **g**, In vitro cardiomyocyte differentiation history tracing (*n* = 2). (top) mESC to cardiomyocyte differentiation timeline and Dam-POI activation. (bottom) Genome-wide clustering of m6A profiles across conditions, like panel f.

For temporal control, Dam is split into two fragments separated by a mutated human estrogen receptor (ER) domain that functions as the intein^40^. Upon addition of 4- hydroxytamoxifen (4OH-TAM), the ER-intein undergoes ligand-dependent protein splicing, reconstituting active Dam. In addition, Dam is tagged with the dTAG degron for rapid proteasomal clearance upon dTAG-13 treatment^41^ (Fig. 1c). We engineered stable mouse embryonic stem cell (mESC) lines expressing these dTAG-Dam-intein-POI. Upon DpnI digestion, which specifically cuts GATC sites bearing m6A^32^, and adapter ligation with high-throughput sequencing, we detected m6A signals only after Dam activation (Fig. 1d, see Methods).

We next benchmarked each Dam-POI (Fig. 1b). Dam-RNAPII^42^ and Dam-Leo1 both correlate strongly with mRNA abundance, serving as transcriptome proxies (Fig. 1d). These factors also accumulate at active enhancers: m6A at enhancer loci correlates with enrichment of H3K27ac, a transcriptionally active histone modification (Fig. 1d). Although Free-Dam (i.e., unfused Dam) is often used to normalize DamID data, its signal also correlates with gene expression because it reflects chromatin accessibility (Extended Data Fig. 1a)^31,32^. Normalizing Dam-RNAPII and Dam-Leo1 signals to Free-Dam did not improve mRNA correlations (Extended Data Fig. 1a), so we omitted this step for those two Dam-POIs. Free-Dam signals in enhancers correlate with ATAC-seq signals, confirming that it can detect accessible enhancers (Fig. 1d).

Dam-Laminb1 marks lamina-associated domains (LADs) in heterochromatin- suppressed regions^35^, showing an inverse pattern relative to Free-Dam and aligning with Laminb1 ChIP-seq (Fig. 1d, Extended Data Fig. 1b). Dam-Taf3^37^ (i.e., two tandem Phd domains) tracks with promoter-associated H3K4me3, reflecting active transcription start sites (Fig. 1d). Dam-H3K27me3 mintbody^37^ (Dam-mH3K27me3) mirrors Polycomb repressive complex 2 (PRC2)-deposited H3K27me3 (Fig. 1d). Finally, based on ChromID’s bivalent-domain profiling strategy^43^, we fused Taf3 Phd domain and Cdh7 chromodomains (Dam-Biv) and observed selective m6A enrichment at regions positive for both H3K4me3 and H3K27me3 (Fig. 1e, Extended Data Fig. 1c).

To assess whether Dam-POI expression perturbs endogenous programs, we performed bulk RNA-seq following activation of Dam-RNAPII and Dam-Laminb1, marking largely nonoverlapping chromatin domains, and found only 28 and 16 differentially expressed genes, respectively (Extended Data Fig. 1d), consistent with prior reports that modest Dam expression minimally impacts cellular physiology^33,42^.

### Whole-genome multimodal history tracing by HisTrac-seq

To demonstrate HisTrac-seq, we performed *in vitro* excitatory neurodifferentiation of mESCs over seven days, which was induced by Doxycycline (Dox)-dependent *Ngn2*- overexpression^44^ (see Methods). By day 2, we observed robust upregulation of the postmitotic neuronal marker *Rbfox3 (NeuN)* and downregulation of the pluripotency marker *Pou5f1* (*Oct4)* (Extended Data Fig. 2a).

We designed three labelling conditions (Fig. 1f): (1) activation of Dam-POIs during the first two days of differentiation, with immediate harvest on day 2 (“Day-2 immature neurons”); (2) activation during the final two days, with harvest on day 7 (“Day-7 mature neurons”); and (3) pulse-chasing, namely transient activation until day 2, followed by harvest on day 7 for retrospective tracing (“Day-2 -> Day7 HisTrac-seq”). Genome-wide m6A quantification and clustering revealed that retrospective HisTrac-seq profiles co-cluster with Day-2 samples rather than Day-7 samples across all seven Dam-POIs (Fig. 1f). Moreover, when we identified stage-specific genes, enhancers, promoters, and genomic bins exhibiting Day-2 or Day-7 enrichment, HisTrac-seq recapitulated the Day- 2 m6A levels (Extended Data Fig. 2b; also see genome-browser tracks in Fig. 1f). Thus, HisTrac-seq can reconstruct epigenetic and transcriptional profiles of the past time point.

To test lineage generality, we differentiated mESCs into cardiomyocytes over seventeen days. By day 10, *Myl7* was induced and *Oct4* was repressed (Extended Data Fig. 2c). Applying Dam-Leo1 and Free-Dam, we compared Day-10, Day-17, and Day-10-> Day-17 HisTrac-seq samples. As with neurons, retrospective labels from Day-10 activation clustered with the Day-10 snapshot, confirming that HisTrac-seq captures past profiles in cardiomyocytes as well (Fig. 2g, also see Extended Data Fig. 2d). Collectively, these data establish HisTrac-seq as a broadly applicable, multimodal history-tracing platform.

**Fig 2.**
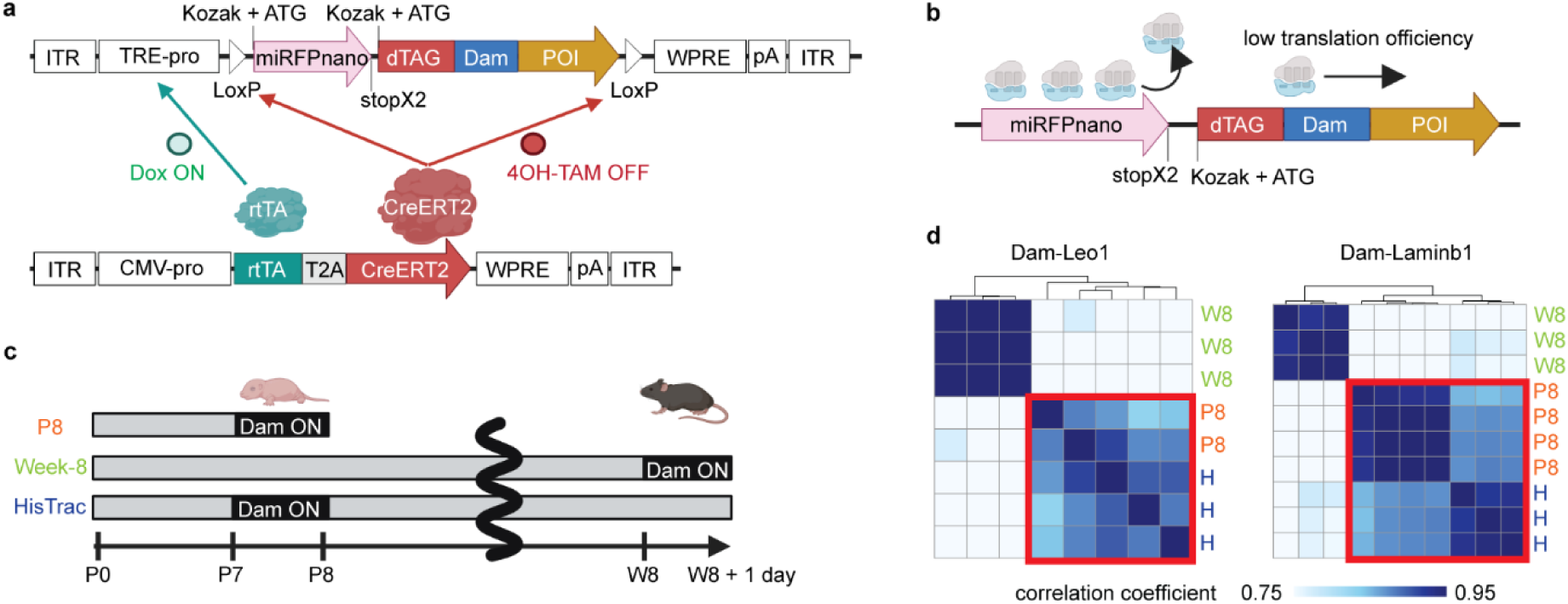
*In vivo* HisTrac-seq. **a**, AAV9 delivery and temporal control of Dam-POI *in vivo*. Dam-POI cassette driven by TRE-Tight2 promoter flanked by LoxP sites; co-delivered rtTA-T2A-CreERT2 under a constitutive CMV promoter. Dox induces Dam-POI expression via rtTA binding, while 4OH-TAM activates CreERT2 to excise the Dam-POI cassette. **b**, Translational reinitiation strategy for tuning Dam-POI expression *in vivo*. A primary open reading frame (ORF) encodes miRFP670nano with a stop codon, followed by the Dam- POI ORF. Ribosomes occasionally reinitiate at the downstream ORF, yielding low-frequency Dam-POI translation that prevents m6A saturation and toxicity. **c**, *In vivo* history-tracing workflow and genome- wide m6A profiling (*n* = 2–4). (top) Activation windows for neonatal (P8), adult (Week-8), and retrospective HisTrac (P8 -> week 8) labelling with Dam-Leo1 and Dam-Laminb1. (bottom) Genome- wide m6A signals quantified in fixed bins and clustered by Pearson’s correlation.

### Whole-genome multimodal history tracing by HisTrac-seq *in vivo*

To implement HisTrac-seq in the mouse brain, we packaged Dam-POI constructs into Adeno-associated virus (AAV) 9 under a tetracycline response element (TRE)-Tight2 promoter flanked by LoxP sites (Fig. 2a). Co-infection with AAV9 expressing rtTA-T2A- CreERT2 enables Dox–dependent induction of Dam-POI and 4OH-TAM-triggered Cre-mediated cassette excision for permanent inactivation. Given variability in injection volume and tissue-sample size, which complicates precise m6A quantification with sequencing, we first validated our Dox-on/4OH-TAM-off system by cortical injection of a control AAV9 vector (EGFP flanked by LoxP sites under the TRE-Tight2 promoter; Extended Data Fig. 3a), followed by EGFP signal quantification on sections. We confirmed that Dox treatment robustly induced EGFP expression, and that 4OH-TAM pretreatment prevented it, demonstrating efficient on/off control (Extended Data Fig. 3b).

Previous studies have shown that harnessing translational reinitiation optimizes Dam–POI expression levels, preventing m6A signal saturation and minimizing the toxic effects of m6A *in vivo*^38,42^. To implement this, we positioned the Dam-POI open reading frame immediately downstream of the miRFP670nano fluorescent-protein ORF (terminated by its stop codon), allowing ribosomes to reinitiate translation of the secondary Dam-POI ORF at low frequency (Fig. 2b).

We then performed transient labelling with Dam-Leo1 (transcription) and Dam- Laminb1 (heterochromatin LADs) in neonatal postnatal day 8 (P8), and adult Week-8, cortex. In each case, we harvested a “snapshot” one day post-activation. For retrospective tracing, we pulse-chased at P7-P8 and harvested at week 8 (P8 -> Week-8 HisTrac-seq) (Fig. 2c). We selected Dam-Leo1 over Dam-RNAPII to obtain the transcriptome because the shorter length of *Leo1* sequence (2 Kb) makes it compatible with AAV packaging over the main subunits of RNAPII (i.e., *Polr2a*, *Polr2b*, 6 Kb and 3.5 Kb). Genome-wide m6A profiles from both Dam-Leo1 and Dam-Lamin B1 show that retrospective samples cluster with the P8 snapshots rather than the adult profiles (Fig. 2c). Stage-specific genes and genomic regions likewise display P8-level m6A in HisTrac- seq samples (Extended Data Fig. 3c). A modest increase in Week 8-specific signal likely reflects residual Dam protein not immediately cleared by Cre excision (Extended Data Fig. 3c); in principle, direct infusion of dTAG-13, which does not cross the blood-brain barrier, could further tighten the labelling window^45^.

Together, these data demonstrate that transient Dam activity can inscribe stable m6A marks that persist on genomic DNA for over two months *in vivo*, enabling reliable, whole-genome history tracing in the brain.

### Tn5 transposase-mediated detection of m6A signals (Dam&Tag)

Previous studies have demonstrated that DamID can be adapted for single-cell sequencing^37,46^. However, current scDamID approaches depend on plate-based library preparation, which limits the number of cells that can be processed in a single experiment and constrains overall throughput. This restriction hampers their application to complex, heterogeneous tissues, such as the brain, that require profiling of large, diverse cell populations. Furthermore, plate-based workflows require specialized equipment and infrastructure, which limits their broader accessibility.

To overcome these limitations, we developed a droplet-based, high-throughput single-cell m6A profiling workflow on the 10x Genomics Chromium platform, a widely adopted system for single-cell genomics. Even though CUT&Tag has been shown to be compatible with the 10x Genomics Chromium scATAC-seq chemistry to map chromatin profiles at single-cell resolution^4,47–50^, we could not find an antibody that robustly recognizes DNA m6A in the chromatin context. To bridge this gap, we repurposed m6A -

Tracer, a catalytically inactive DpnI variant previously used for live-cell m6A imaging^35^. We fused it to a V5 epitope and EGFP tags (V5-EGFP-m6A-Tracer; Fig. 3a). After purification of the tracer protein (Methods), we applied it to permeabilized nuclei, then targeted Protein A-Tn5 transposase to the m6A-marked sites by incubating with anti-V5 or anti-EGFP antibodies. This “Dam&Tag” protocol enables Tn5-mediated sequencing adapter insertion (i.e., tagmentation) specifically at m6A-modified regions (Fig. 3a).

**Fig 3.**
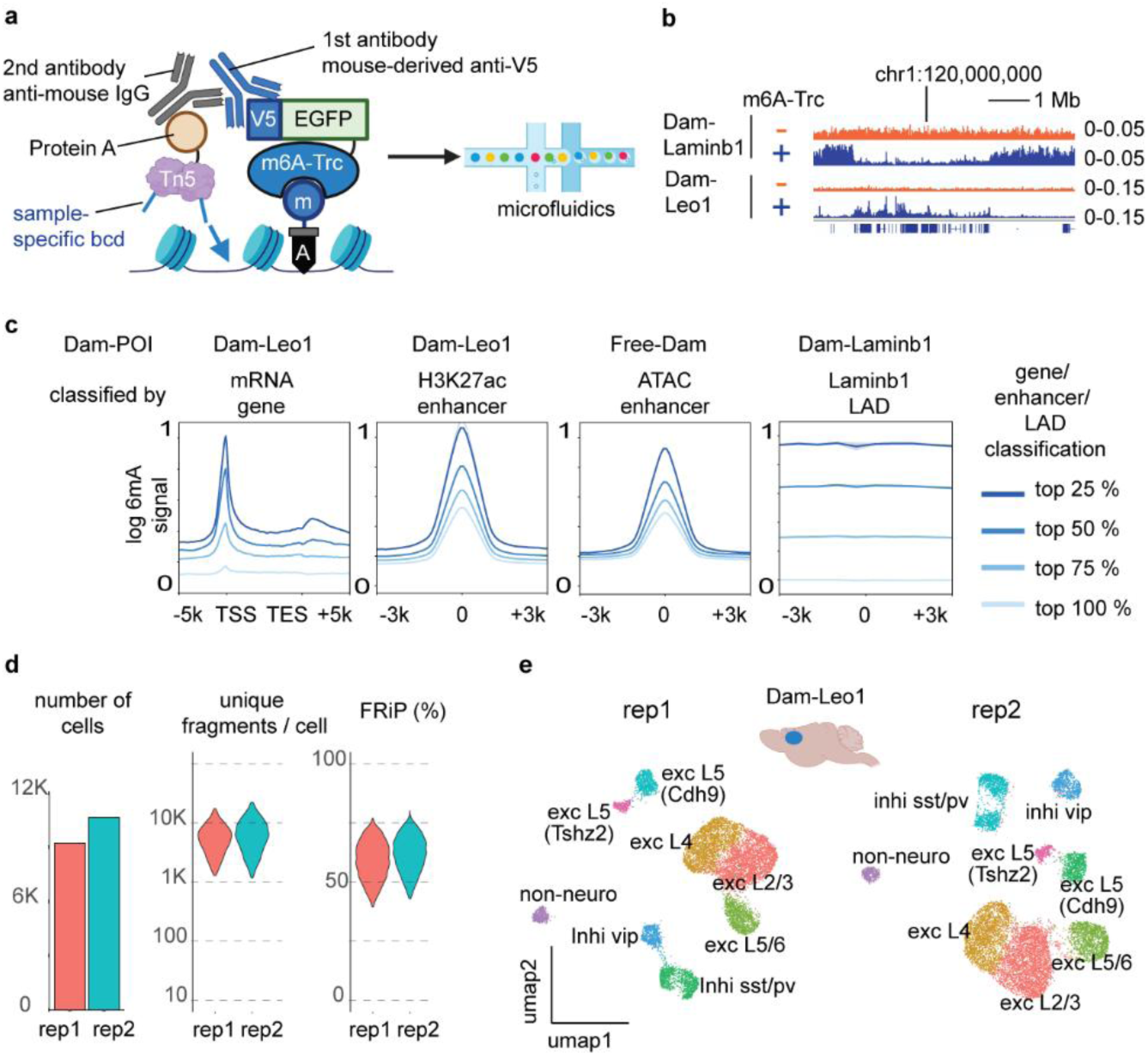
High-throughput single-cell m6A mapping by scDam&Tag. **a**, Workflow of scDam&Tag. Permeabilized nuclei are incubated with V5-EGFP-m6A-Tracer (m6A-Trc) to recognize genomic m6A. An anti-V5 antibody recruits Protein A–Tn5 transposase loaded with ME-A adapters bearing sample-specific barcodes (bcd). Nuclei undergo single-cell library preparation on the microfluidics system. **b**, Genome browser tracks of m6A signals in mESCs with bulk Dam&Tag using Dam-Leo1 and Dam-Laminb1. Comparisons between m6A-Tracer-negative (orange) and tracer-positive (blue) conditions confirm that POI-specific signals can be detected only in the m6A-Tracer-positive condition. Dam-Laminb1 marks heterochromatin in regions mutually exclusive with Dam-Leo1, without Free-Dam normalization (*n* = 3). **c**, Meta-analysis of bulk Dam&Tag specificity in mESCs (*n* = 2). Genomic features (genes, enhancers, and LADs) are stratified into quartiles (25%, 50%, 75%, 100%) by activity (mRNA-seq, H3K27ac-ChIP, ATAC-seq, Laminb1-ChIP). Plots show mean m6A enrichment ± s.e.m. **d**, Quality control metrics for cortical neuron scDam&Tag (Dam-Leo1) (*n* = 2). (left) Number of nuclei recovered per biological replicate. (middle) Violin plots showing unique fragment counts per cell. (right) Violin plots showing FrIP per cell. **e**, UMAP embedding per replicate with cell type labels. Exc, excitatory neurons; inhi, inhibitory neurons; L, cortical layer.

We screened seven commercial V5 and EGFP antibodies by comparing bulk library yields from Dam-Leo1 and Dam-Laminb1 mESCs with and without V5-EGFP-m6A- Tracer (Extended Data Fig. 4a). Five antibodies showed tracer-dependent signals (Extended Data Fig. 4b), and among these, sequencing revealed that the mouse monoclonal anti-V5 clone SV5-pk1 produced the lowest background tagmentation (Fig. 3b, Extended Data Fig. 4c, see Methods).

Using SV5-pk1–based Dam&Tag, we profiled Dam-Leo1, Free-Dam, and Dam- Laminb1 in bulk cells and observed the expected positive correlations with mRNA levels, H3K27ac/enhancer accessibility, and Laminb1 ChIP-seq, respectively (Fig. 3c), mirroring our DpnI-digestion results. Finally, applying Dam&Tag to our *in vitro* neurodifferentiation model confirmed that this protocol reliably captures retrospective chromatin signatures (Extended Data Fig. 4d).

### Single-cell m6A signal detection with scDam&Tag

We then established scDam&Tag (workflow in Extended Data Fig. 5a) by combining two key innovations. First, we adopted the nanobody-mediated CUT&Tag (nanoCT) two-step tagmentation protocol to maximize library complexity: an initial Tn5 reaction inserts ME-A adapters, followed by a second round of ME-B adapter insertion after the linear amplification step of 10x Chromium microfluidics chemistry^4^. Second, to reduce per-sample cost and batch effects, we applied a combinatorial-indexing scheme inspired by previous studies (e.g., SUM-seq^51^). In this design, the first tagmentation uses ME-A adapters carrying sample-specific barcodes; after pooling, overloaded nuclei enter the 10x Chromium microfluidic device in droplets containing multiple nuclei. Inside each droplet, adapters are further indexed by cell-specific barcodes. This two-tier indexing retains both the sample origin and single-cell identity of every read, enabling high-throughput, multiplexed m6A profiling within a single Chromium channel.

For proof of principle, we applied scDam&Tag to adult mouse cortex expressing Dam-Leo1 via AAV9 (Fig. 3d,e, Extended Data Fig. 5a). After tracer-mediated tagmentation, we enriched m6A-marked neuronal nuclei by FACS based on EGFP signal from V5-EGFP-m6A-Tracer (Extended Data Fig. 5b). We pooled two biological replicates for library preparation and sequencing. Post-demultiplexing and quality control, we recovered 9,232 and 10,656 cells, with median unique fragment counts of 5,264 and 5,947, and Fractions of Reads in Peaks (FrIP) of 59 % and 62 % (Fig. 3d, Supplementary Table 1). The complexity metric ranks among the highest for droplet-based single-cell chromatin assays using similar microfluidics platforms (e.g., scCUT&Tag^47,50^, nanoCT^4^, NTT-seq^49^, scPaired-Tag^48^).

Clustering and uniform manifold approximation and projection (UMAP) revealed that over 95 % of captured cells were neurons, reflecting AAV9’s tropism, and within this population, we delineated major cortical excitatory and inhibitory subtypes (Fig. 3e, Extended Data Fig. 5c).

In summary, scDam&Tag achieves both high cell-number throughput and high library complexity, enabling comprehensive *in vivo* profiling of diverse cell types. Moreover, as will be demonstrated below, by multiplexing up to six biological samples in a single 10X Chromium controller channel, we have achieved recovery of as many as 36,805 cells (Fig. 4; Supplementary Table 1), dramatically reducing per-cell and per-sample library costs.

**Fig 4.**
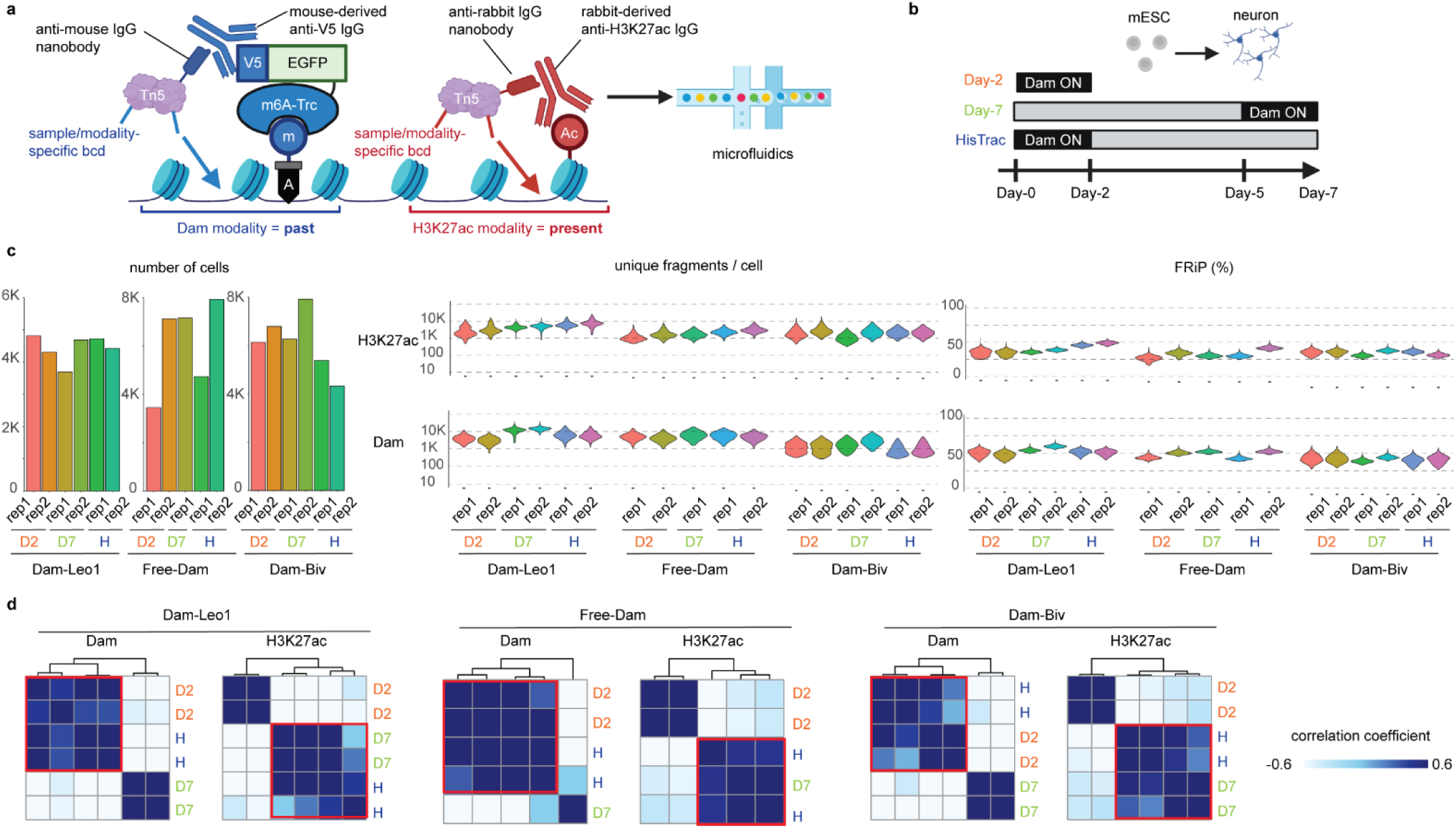
Single-cell HisTrac-seq simultaneously profiles the past and present. **a**, Dual-CUT&Tag workflow of scHisTrac-seq. m6A-Trc binds genomic m6A marks (historical mark), which are recognized by a mouse-derived anti-V5 IgG and tagged with anti-mouse-IgG nanobody-Tn5. Present transcriptional activity is profiled in parallel using rabbit-derived anti-H3K27ac IgG and anti-rabbit-IgG nanobody-Tn5. Modality- and sample-specific barcodes are incorporated via ME-A adapters. Libraries are then prepared on microfluidics platform. **b**, Timeline of mESC neurodifferentiation and Dam-POI activation windows. **c**, Quality control metrics for snapshot scDam&Tag (Day-2, Day-7) and scHisTrac-seq. (left) Number of nuclei recovered per biological replicate. (middle) Violin plots showing unique fragment counts per cell. (right) Violin plots showing FrIP per cell. **d**, Pseudo-bulk analysis of sample clustering for each modality. Color-coded by Pearson’s correlation. The sample size is indicated in panel **c**.

### Single-cell history tracing with scHisTrac-seq

We next extended scDam&Tag into scHisTrac-seq to capture both past and present molecular profiles in the same single cells. Leveraging the nanobody-Tn5 transposase-based approaches of nanoCT and NNT-seq^4,49^, we performed a dual CUT&Tag reaction: m6A marks (historical signal) are recognized by mouse-derived anti- V5 antibody and tagged with anti-mouse-IgG nanobody-Tn5 transposase, while the present molecular profile (i.e., transcriptome) is detected by rabbit-derived anti-H3K27ac antibody and anti-rabbit-IgG nanobody-Tn5 transposase. H3K27ac reflects transcriptional activity of genes and enhancers. Modality-specific barcodes embedded in the ME-A adapters allow retrospective demultiplexing of the two data streams after sequencing (Fig. 4a; Extended Data Fig. 6a).

Using mESC-derived excitatory neurons, we applied scHisTrac-seq with three Dam-POIs, namely, Dam-Leo1, Free-Dam, and Dam-Biv, across three conditions (Day- 2, Day-7, and Day-2 -> Day-7 HisTrac, Fig. 4b). Six biological samples (two replicates per condition) were pooled into a single 10x Chromium channel via sample-indexed ME-A adapters (Extended Data Fig. 6; Supplementary Table 1). After quality control, we recovered 3,450–7,475 cells per sample, and in total 93,844 cells (26,648 for Dam-Leo1, 30,391 for Free-Dam, and 36.805 for Dam-Biv). Median unique fragment counts per cell ranged from 2,948-14,330 for Dam-Leo1, 3,985-5,974 for Free-Dam, 980-3,682 for Dam- Biv, and 1,231-7,413 for H3K27ac. Median FrIP was 47-59 % (Dam-Leo1), 42-52 % (Free-Dam), 42-47 % (Dam-Biv), and 28-43 % (H3K27ac) (Fig. 4c; Supplementary Table 1). The lower complexity of Dam-Biv reflects the limited genomic span of bivalent domains.

In pseudo bulk analyses, scHisTrac-seq profiles clustered with Day-7 in the H3K27ac modality but with Day-2 in all Dam modalities, confirming that H3K27ac reports the present profiles while Dam records past (Fig. 4d).

In addition, to assess Tn5 barcode hopping after pooling tagmented nuclei^51^, we examined whether cells bearing Day-2 versus Day-7 sample barcodes would cluster separately. Indeed, cells segregated cleanly into two clusters corresponding to their Day- 2 or Day-7 barcodes (Extended Data Fig. 7; also see Fig. 5a). We reasoned that any barcode hopping would create discordance between a cell’s sample-barcode assignment and its post hoc clustering-based identity. Yet only 0.2 % of Dam-Leo1 cells, 1.0 % of Free-Dam cells, and 0.4 % of Dam-Biv cells exhibited such mismatches, demonstrating that barcode hopping is effectively negligible (Extended Data Fig. 7).

**Fig 5.**
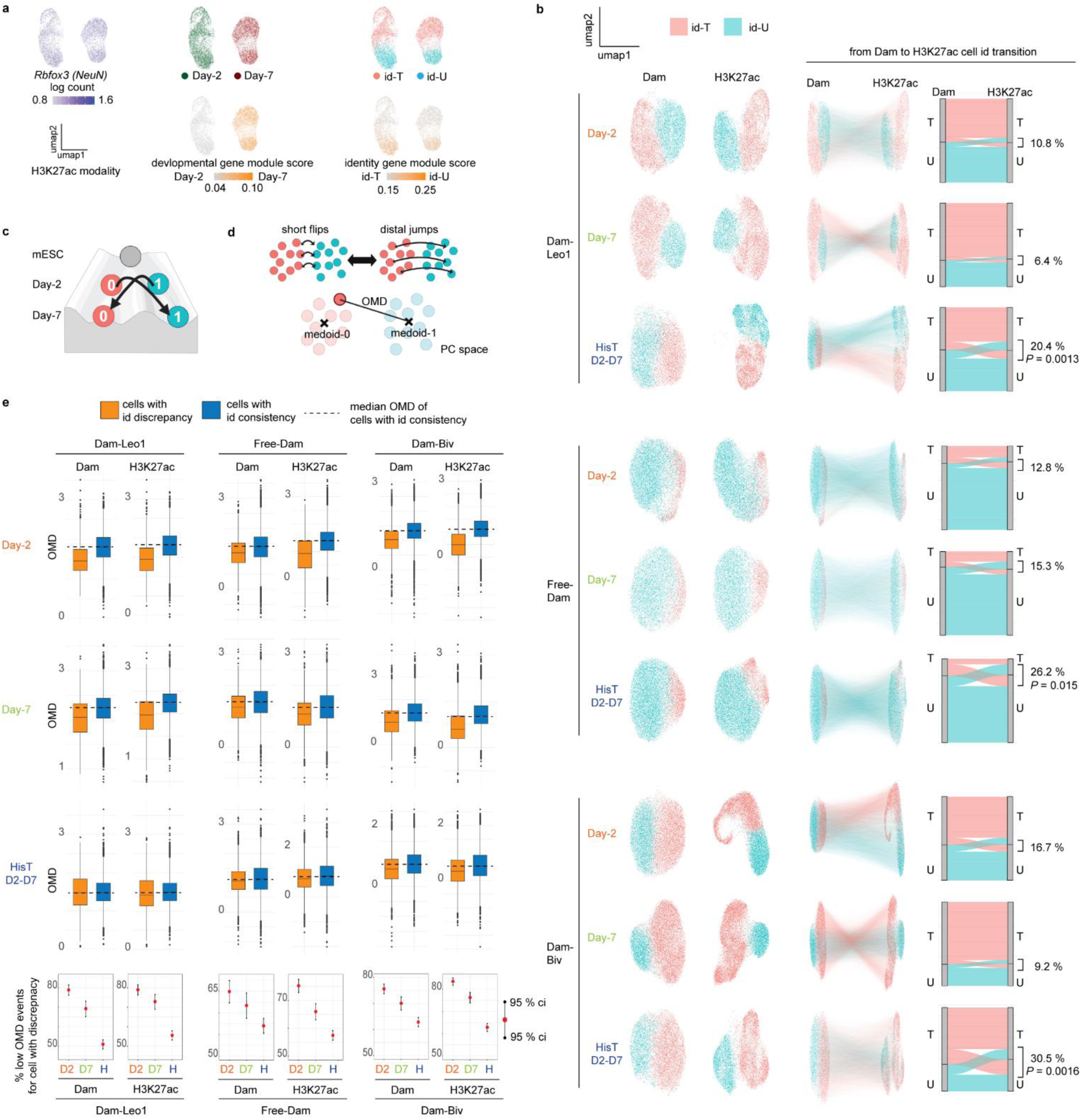
Analysis of cell identity transition during development by scHisTrac-seq. **a**, UMAP embedding of H3K27ac modality from Dam-Leo1 samples at Day-2 and Day-7. Cells primarily cluster by developmental stages and are further subdivided into two identities, namely id-*Tshz2* (id-T) vs. id-*Unc5c* (id-U). They are all *Rbfox3* (*NeuN*)-positive postmitotic neurons. Developmental and identity gene module scores are also shown (bottom). *n* = 2. **b**, (left) UMAP embedding of individual Dam-POIs (Dam-Leo1 – 26,748 cells, Free-Dam - 30,391 cells, Dam-Biv - 36,805 cells), two modalities (Dam, H3K27ac), and three conditions (Day-2, Day-7, HisTrac). Colour denotes id-T and id-U. (right) Analysis of cell identity transition from Dam to H3K27ac modalities. Colour code is based on Dam-modality, and the lines connect the same single-cells across two modalities. Alluvial plots quantify cell identity transition, highlighting the discrepancies between the two modalities. *P* values are calculated by one-sided Welch’s two-sample *t*-tests. The sample size is indicated in Fig. 4c. **c**, Scheme representing “identity jumps” from one canal to another in Waddington landscape. **d**. Definition of opposing medoid distance (OMD) to distinguish short boundary flips from long-distance jumps of identities on the PC space. **e**. Quantifying identity ambiguity with OMD. (top) Boxplots of OMD distribution on Dam and H3K27ac PC spaces for individual Dam-POI and conditions. Comparing cells showing consistent identities (blue) versus discrepant identities(orange) between two modalities. (bottom) Among cells with identity discrepancy, the fraction of cells showing OMD falls below the median OMD of consistent cells (dashed line in boxplots) is shown as red dots with 95 % confidence intervals. HisTrac samples exhibit fewer low-OMD events than snapshot conditions, indicating true long-distance identity jumps. The sample size is indicated in Fig. 4c.

### Characterization of disruptive cell identity jumps during development

We next investigated developmental transitions, using scHisTrac-seq data. Clustering of each condition, namely, Day-2, Day-7, and HisTrac, revealed two reproducible subpopulations across all Dam-POI and H3K27ac modalities at both Day-2 and Day-7 stages (Fig. 5a,b, Extended Data Fig. 8a).

On top of 1,392 Day-2 vs. Day-7 stage-specific genes (developmental gene module in Fig. 5a, Extended Data Fig. 8a), we identified 449 identity-specific genes consistently differentially expressed between subpopulations at both Day-2 and Day-7 (identity gene module in Fig. 5a, Extended Data Fig. 8a) (Supplementary Table 2). Gene- ontology analysis revealed that identity genes are enriched for neuronal projection, sensory system development, and intracellular signals (Extended Data Fig. 8b). Because identity-specific genes include *Tshz-2* and *Unc5c,* which show mutually exclusive expression patterns in cortical and basolateral amygdala excitatory neurons (Extended Data Fig. 8a), we named these subpopulations as id-*Tshz2* (id-T) and id-*Unc5c* (id-U).

Because traditional snapshot data (i.e., Day-2 and Day-7) clusters primarily by developmental stage, it is very challenging to infer single-cell trajectories between Day-2 and Day-7, obscuring transitions between id-T and id-U (Fig. 5a). Leveraging scHisTrac- seq’s dual modalities, Dam marking past (Day-2) and H3K27ac reporting present (Day- 7), we tracked each cell’s identity over time. Most cells maintained their initial identity, but a substantial fraction displayed discrepancies: 20.4 % (Dam-Leo1), 26.2 % (Free-Dam), and 30.5 % (Dam-Biv) of cells switched identity from Day-2 to Day-7 (Fig. 5b). These rates significantly exceed the background discrepancies in independent Day-2 and Day- 7 snapshots (Fig. 5b; see Extended Data Fig. 9 for individual replicates), indicating bona fide disruptive “identity jumps” during differentiation (Fig. 5c).

To distinguish true identity jumps (i.e., long-range transitions) from short-range boundary-level flips between cell clusters (which may result from technical noise, such as nuclear multiplets in droplet reaction), we devised the Opposing Medoid Distance (OMD) metric (Fig. 5d). For each cell, OMD quantifies its distance in principal component (PC) space to the medoid of the alternate identity cluster. Accordingly, a low OMD indicates that a cell lies close to the alternate-cluster medoid, reflecting an ambiguous identity. We quantified OMD for individual Dam-POI samples, modalities, and conditions. In snapshot data, cells with discordant identities between Dam and H3K27ac modalities exhibited lower OMD than those with consistent identities on both Dam and H3K27ac modalities (Fig. 5e), confirming that these apparent mismatches stem from ambiguous boundary cases rather than true transitions. In contrast, in HisTrac, such trends were significantly attenuated (Fig. 5e), demonstrating that these are true, disruptive switches between id-T and id-U from Day-2 to Day-7.

### Epigenetic and transcriptional mechanisms of identity jumps

We next dissected the molecular underpinnings of identity jumps. To uncover transcriptional drivers of identity jumps, we compared gene expression in cells undergoing jumps against those maintaining identity using the HisTrac Dam-Leo1 modality. This analysis identified 162 jump-associated genes for id-U -> id-T transition, while only 13 genes for id-T -> id-U transition (Fig. 6a, Extended Data Fig. 10a, Supplemental Table 2). We mainly focused our downstream analyses on the id-U -> id-T transition. Notably, the 162 jump-associated genes are normally upregulated at Day-7 relative to Day-2, indicating that premature activation of portions of the differentiation program accompanies these identity switches (Fig. 6a, also see Fig. 10a for id-T -> id-U jump showing the same trend). Gene ontology enrichment highlights roles in intracellular communication, membrane transport, and extracellular matrix organization, suggesting that altered cell-environment interactions may precipitate identity jumps (Fig. 6b).

**Fig 6.**
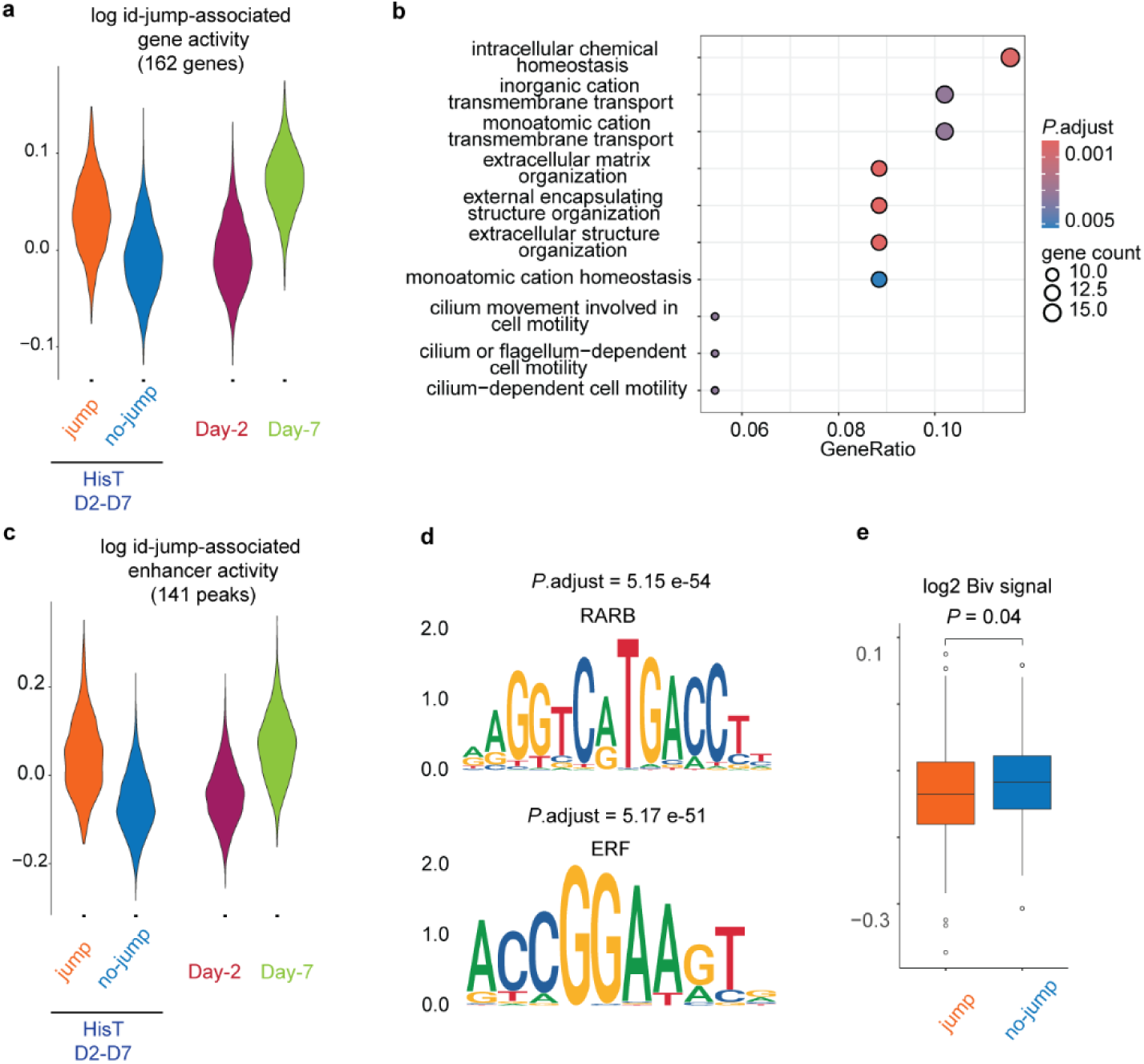
Epigenetic and transcriptional basis of identity jumps. **a**, Violin plots of Dam-Leo1-measured expression for the 162 genes associated with identity jumps adjusted *P*-value < 0.05). Cells that undergo jumps (orange) versus those that maintain identity blue) are compared. These genes normally show higher expression at Day-7 (green) versus Day-2 dark red) (*n* = 2). **b**, Gene ontology analysis of 162 jump-associated genes. **c**, Violin plots of Dam-Leo1 signal at the 141 peaks associated with identity jumps (adjusted *P*-value < 0.05). Jumping cells orange) versus non-jumping cells (blue) are compared. These enhancers are typically more active at Day-7 (green) than Day-2 (dark red) (*n* = 2). **d**, Transcription factor–motif enrichment analysis in the 141 jump-associated enhancers, with RARB and ERF motifs among the most significantly enriched. **e**, Boxplots of normalized Dam-Biv (bivalency) signal at jump-associated 141 enhancers, comparing umping versus non-jumping cells (*n* = 2). A loss of bivalency accompanies identity jumps. *P*-value by Welch’s two-sample, two-sided *t*-test, t = -2.003, df = 268.96.

We next examined the chromatin basis of these transitions. Screening for differential Dam-Leo1 revealed 141 peaks associated with id-U -> id-T (Fig. 6c, Supplementary Table 2), all of which are distal enhancers (Extended Data Fig. 10b). Like the jump-associated genes, these enhancers exhibit higher activity at Day-7 as compared to Day-2, consistent with their precocious engagement in jumping cells (Fig. 6c). Transcription factor motif analysis of these enhancers uncovered enrichment for Retinoic Acid Receptor Beta (RARB) and Ets Receptor Factor (ERF) binding sites (Fig. 6d, Extended Data Fig. 10, Supplementary Table). In the developing cortex, RARβ, often paired with RXRγ, is enriched anteriorly, where it governs area-specific thalamocortical connectivity, dendritic spinogenesis, and layer-4 marker RORβ expression^52^. ERF, induced by retinoic acid, is critical for neurogenesis in forebrain progenitors, antagonizing ETS-driven proliferative cues to permit neuronal differentiation^53^. These pathways likely orchestrate the enhancer activation that drives identity jumps.

Finally, we assessed the role of bivalent chromatin in maintaining developmental plasticity. Bivalency regulates plasticity of gene inducibility by keeping genes in a poised state, marked simultaneously by H3K4me3 and H3K27me3, which allows for rapid and reversible transcriptional switching in response to differentiation signals^39^. Using pseudo bulk profiles from the Dam-Biv and Free-Dam modalities with normalization of Dam-Biv by Free-Dam, we quantified bivalency at the 141 jump-associated enhancers. Jumping cells exhibited a loss of bivalency (i.e., resolution from the poised H3K4me3/H3K27me3 dual state to active chromatin) at these sites (Fig. 6e). This resolution of enhancer bivalency appears to underlie the disruptive identity transitions observed during differentiation.

Together, HisTrac-seq provides the first single-cell and whole-genome demonstration of disruptive cell identity jumps during development. These jumps are underpinned by premature activation of maturation programs, spanning cell-environment interactions, morphogen to transcription factors signalling, enhancer activation, and epigenetic remodelling. By revealing this hidden layer of developmental plasticity, scHisTrac-seq opens new avenues for dissecting how abrupt identity transitions shape tissue formation, function, and disease.

## DISCUSSION

It remains a major challenge in genomics to move beyond static snapshots and capture truly time-resolved information. Although DNA-recording methods^9–12,14–28^ that inscribe event history into the genome have advanced rapidly (e.g., DNA Typewriter- ENGRAM^12^, DCM-TM^14^), multi-time-point whole-genome observation at single-cell resolution remains severely limited. Live-seq provides an orthogonal solution by physically extracting RNA from living cells but requires continuous optical and mechanical access to individual cells, severely limiting cell number-throughput^13^. Furthermore, these technologies provide only one modality, typically the transcriptome. HisTrac-seq overcomes these barriers by enabling simultaneous, single-cell resolution profiling of both the present and a defined past time point across the entire transcriptome and epigenome. By leveraging the widely adopted 10x Genomics Chromium platform, we can trace the history of tens of thousands of individual cells in parallel. HisTrac-seq is a broadly applicable “temporal multi-omics” platform to address diverse biological questions from development and learning to disease progression, in versatile experimental systems, since DamID has been validated across a variety of cell types and organisms (summarized in Extended Data Fig. 11).

In this study, we analysed neuronal identity transitions in over 93,000 single cells across multiple epigenetic and transcriptional modalities, reliably revealing disruptive “identity jumps” (schematically, cell transitions from one canal to another on the Waddington landscape, see Fig. 5c). Conventional snapshot-based inference methods cannot resolve them due to the requirement of continuity among single-cell snapshots. Furthermore, we dissected the molecular underpinnings of jumps, including temporal shifts in gene expression, enhancer activation, bivalent marking, environmental cues, and signalling-to–transcription factor cascades. These findings provide a new insight into how to distinguish stable cell types and transient cell states. By quantifying transition probabilities, HisTrac-seq offers a framework for mapping an “energy landscape” of fate decisions during development. While current work was carried out in a simple monoculture of neurons, future work may explore how experimental manipulations (e.g., morphogen, epigenetic chromatin remodeling, cell-cell interactions, three-dimensional culture, *in vivo* contexts, or disease conditions) reshape this landscape. This will allow us to understand how the height of energy barriers between cell fates is determined.

HisTrac-seq does have limitations. First, it records only two time points. Extending to three or more would require orthogonal DNA marks (e.g., DCM) or chemical labels (e.g., biotin), or integration with CRISPR-based sequential recorders: combining HisTrac’s genome-wide scope with CRISPR’s finer temporal resolution^12^ could yield a comprehensive dynamic readout. Second, because m6A marks are diluted with DNA replication. Neurons give rise to diverse cell types and carry out complex computational functions as postmitotic cells, and as we have shown, non-neuronal postmitotic cells (e.g., cardiomyocytes) are also compatible with long-term tracing. However, engineering m6A- maintenance machinery would further extend this method’s applicability to proliferative lineages. Dam variants that recognize hemi-methylated GATC sites, as previously proposed^54^, may be intriguing tools. Furthermore, a recent preprint proposed that m6A in replicated DNA may be detected with an anti-m6A antibody following genomic DNA denaturation, potentially allowing history tracing after one round of DNA replication^55^. Third, although droplet-based Chromium workflows deliver high cell counts, they trade depth per cell and demand substantial input material. For rare populations requiring deeper sequencing coverage, plate-based scDamID^37^ may remain preferable and may also reduce background from Tn5-mediated open-chromatin labelling.

Finally, the V5-EGFP-m6A-Tracer and Tn5-dependent Dam&Tag approach opens exciting avenues for “spatiotemporal multi-omics” by combining with emerging spatial omics technologies. HisTrac-seq should integrate seamlessly with spatial chromatin platforms (e.g., DBiT-seq^56^, Slide-tag^57^) that deploy Tn5 for histone or accessibility mapping, enabling concurrent profiling of past and present profiles within intact tissue architecture. Moreover, because m6A-Tracer is amenable to fluorescence imaging^35^, combining EGFP readouts with high-resolution FISH (e.g., DNA seqFISH^58^) could visualize the historical and present regulatory status of individual loci while preserving subnuclear context. Together, these advances position HisTrac-seq as a versatile toolkit for unravelling dynamic biological processes in space and time.

## Methods

### Vector construction

For piggyBac-based Dam-POI overexpression in mESCs, a piggyBac backbone vector containing a gene cassette flanked by piggyBac terminal inverted repeats (ITRs) was used. The cassette comprised the CAG promoter*, FKBP12^F36V^-Dam-intein-SwaI* restriction site, followed by an internal ribosome entry site (*IRES*) driving expression of a blasticidin resistance gene and a polyadenylation signal. DNA fragments encoding a protein of interest (POI) were amplified by PCR using plasmids obtained from Addgene as templates and inserted into the SwaI-digested vector using NEBuilder HiFi DNA Assembly (New England Biolabs, E2621). The vector lacking a POI insert (Free-Dam- intein) was used as a control. *Dam-intein* was clone from *p-attB-FL.hsp70P-Dam[4-HT- intein@L127C]Myc[open]*^40^ (Addgene plasmid # 71805, a gift from Bas van Steensel). *Leo1* was cloned from *pcDNA3 Leo1*^59^ (Addgene plasmid # 11061, a gift from Matthew Meyerson), *RNAPII* was cloned from *FLAG-Pol-II-WT*^60^ (Addgene plasmid # 35175, a gift from Benjamin Blencowe), *Laminb1* was cloned from *pCIBN-EYFP-hLaminB1*^61^ (Addgene plasmid # 103800, a gift from Karsten Rippe). Anti-H3K27me3-mintbody was cloned from *PB533_2E12LI-sfGFP*^62^ (Addgene plasmid # 167527, a gift from Hiroshi Kimura). *2xTaf3* phd domain, and *Biv* (i.e., *Taf3* phd domain + *Cdh7* chromodomain) were cloned from *pv9_TAF3_2xPHD-13XL-BASU and pv9_CBX7_Chromo/TAF3.PHD-13XL- BASU*^43^ (Addgene plasmid # 174914 and 174915, gifts from Tuncay Baubec).

For AAV-based DamID, the cassette comprised a doxycycline-inducible *TRE Tight2* promoter, followed downstream by a *loxP* site, *miRFPnano* with two stop codons, *Kozak-ATG-FKBP12^F36V^ -Dam*, a SwaI restriction site for inserting a protein of interest (POI), a second *loxP* site, hepatitis virus posttranscriptional regulatory element (WPRE), and a polyadenylation signal. DNA fragments encoding POIs were obtained either by PCR using template plasmids obtained from Addgene and assembled into the SwaI-digested vector using NEBuilder HiFi DNA Assembly. *CMV-Promoter::rtTA-T2A-CreERT2* was synthesized by VectorBuilder.

AAV packaging was performed by the Viral Vector Facility (VVF, Zurich, Switzerland)

For V5-EGFP-m6A-Tracer protein puroductin, *EGFP-m6A-Tracer* was cloned from *m6A- Tracer-NES*^63^ (Addgene plasmid # 159607, a gift from Aaron Streets) and inserted into *pTXB1* Vector (NEB, N6707; for IMPACT system) with *T7* tag and *V5* tag using NEBuilder HiFi DNA Assembly.

Dam-POI overexpressing plasmids and *pTXB1-V5-EGFP-m6A-Tracer* plasmid will be deposited on Addgene.

### Protein production

Tn5 transposase was generated from *pTXB1-Tn5*^64^ (Addgene plasmid # 60240, a gift from Rickard Sandberg). Anti-mouse IgG nanobody- and anti-rabbit IgG nanobody-fused Tn5 transposases were generated from *pTXB1-nbMmKappa-Tn5* and *pTXB1-nbOcIgG- Tn5*^49^ (Addgene plasmid # 184286 and 184285, gifts from New York Genome Center & Ivan Raimondi). V5-EGFP-m6A-Tracer was generated from *pTXB1-V5-EGFP-m6A- Tracer* (described above). We used the NEB IMPACT system following previous studies^49,64^. T7 Express Competent E. coli (NEB, C2566) was transformed with pTXB1 plasmds and a single colony is used to inoculate 5 mL LB/Amp overnight; this starter culture (5 mL) seeds 1 L LB/Amp, grown at 37 °C to around A₆₀₀ = 1.0, chilled, induced with 0.25 mM IPTG, and grown at 23 °C to A₆₀₀ = 2.0 before harvesting by centrifugation and snap freezing. Cell pellets are lysed in HEGX buffer (20 mM HEPES pH 7.5, 0.8 M NaCl, 1 mM EDTA, 10% glycerol, 0.2% Triton x-100) with 1x protease inhibitors (Sigma, 11836170001) by sonication, clarified at 14,000x g, treated with 10 % PEI to remove nucleic acids, and the resulting supernatant is applied to a chitin affinity column (NEB, S6651) at 4 °C; after washing, the column is charged with 100mM DTT-containing HEGX buffer supplimented with protease inhibtors and incubated 48 h at 4 °C for intein-mediated cleavage, then 1 mL fractions are eluted and assayed by Coomassie Blue staining before pooling and dialysis. Dialysis was perfromed in 2x Dialysis buffer (100mM HEPES pH7.5, 0.2 M NaCl, 0.2 mM EDTA, 20% glycerol, 2.0 mM DTT, 0.2% Triton x-100) with 3.5 KDa CelluSep T1 Tubings (Spectrum Laboratories, SP3 #132720) for 48 hours at 4 °C. Purified proteins are stored in 50% glycerol at -80 °C for long-term and at -20 °C for short-term.

### Adapter loading to Tn5 transposase

For Tn5 transposase and Protein A -Tn5 transposase loaded with conventional ME-A/ME- B mixed adapters, ME-A + ME-rev and ME-B + ME-rev oligos (Supplementary Table 3) were used. For Tn5 transposase loaded only with ME-B adapter, ME-B + ME-rev oligos were used. For Protein A -Tn5 transposase, Mouse nanobody-Tn5 transposase, and Rabbit nanobody-Tn5 transposase, loaded with sample/modality-barcoded ME-A adapters, ME-A-index-series + ME-rev oligos (Supplementary Table 3) were used. ME- A-index-series oligos were designed based on NTT-seq^49^ and nanoCT^4,65^. 100 µM adapter mixtures in 10 mM Tris pH 7.5 were denatured at 95 °C for 5 minutes and gradually cooled (5 °C every 3 minutes in a thermalcycler) down to room temperature. 14.5 µM transposases were mixed with 1/8x volume of adapter mix, incubated at room temperature, and stored at -20 °C. We optimized the concentration of ME-B-only Tn5 transposase for tagmentation of linear amplified DNA following the previous study^65^.

### mESC culture and stable cell line preparation

mESCs harboring a Dox-inducible *Ngn2* (*Ngn2*-mESCs) are a kind gift from Dirk Schubeler (Basel, Switzerland)^66^. They were maintained in ESC medium, namely DMEM (Thermofisher, 41966029) supplemented with 15% fetal bovine serum (Thermofisher, 10437-028), 1× GlutaMax (Thermofisher, 35050061), 1× nonessential amino acids (Thermofisher, 11140035), 110 µM beta-mercaptoethanol (Thermofisher, 21985023), 1x Penicillin-Streptomycin (Thermofisher, 15140122), and leukemia inhibitory factor (LIF; Miltenyi Biotec, 130-095-779) on plates coated with 0.1% gelatin (Sigma). A piggyBac- DamIntein-POI plasmids were co-transfected with a plasmid encoding piggyBac transposase (pBase) using Lipofectamine 2000 (Thermofisher, 11668019). Starting the following day, 500 µg mL⁻¹ G418 (Thermofisher, 10131035) and 10 µg mL⁻¹ Blasticidin (Thermofisher, 11583677) were added to the culture medium to select for stable integration of the piggyBac cassette. Selection was continued for 2 weeks, and we picked up resistant clones. For each stable cell line clone, we confirmed the induction of Dam-POI with Dam-qPCR^67^ and DamID-sequencing.

### *Ngn2*-mESC neurodifferentiation and Dam activation

Culture plates were pre-coated with poly-D-lysine (Sigma, P-7886) and laminin (Sigma, L2020). After passage (day 0), *Ngn2*-mESCs were cultured for two days in 1^st^- Neuromedium consisting of DMEM/F12 (Thermofisher, 11320033), 1× GlutaMax, 1× B-27 Supplement minus vitamin A (Thermofisher, 12587010), 1× N-2 Supplement (Thermofisher, 17502048), and 1× Penicillin-Streptomycin, supplemented with 1 µg/mL Dox to induce *Ngn2* overexpression. To promote further neuronal maturation, the medium was replaced by 2^nd^-Neuromedium composed of Neurobasal medium (Thermofisher, 21103049), 1x B-27 Supplement with vitamin A (Thermofisher, 17504044), 1x N-2 Supplement, and Penicillin-Streptomycin, supplemented with 10 ng/mL each of BDNF (PeproTech, 450-02), GDNF (PeproTech, 450-10), and NT-3 (PeproTech, 450-03) for five days. For HisTrac-seq experiments, *Ngn2*-mESCs were pretreated with 1 µM dTAG-13 (Tocris, 6605) for 48 hours to degrade pre-existing DamPOIs in ESC medium. Dam activation was induced by 1 µM 4OH-TAM (Sigma, H6278) for the first two days in 1^st^- Neuromedium. Subsequently, cells were washed multiple times to thoroughly remove residual 4OH-TAM and cultured with 5 µM dTAG-13 to degrade Dam protein until day 7. For the Day-7 snapshot condition, 4-OHT was applied on days 6–7.

### Cardiomyocyte differentiation and Dam activation

Culture plates were pre-coated with Geltrex (Thermofisher, A1413302). We used the PSC Cardiomyocyte Differentiation Kit (Thermofisher, A2921201). After leaching 50% confluency in ESC medium, the medium was changed to Differentiation Medium-A (Thermofisher, A29209-01) (day 0). After 48 hours, the medium was cultured in Differentiation Medium-B (Thermofisher, A29210-01) for 48 hours, and the medium was subsequently replaced by Caridomyocyte Maintenance Medium (Thermofisher, A29208- 01). Medium was replaced every other day. For HisTrac-seq experiments, cells were pretreated with 1 µM dTAG-13 (Tocris, 6605) for 48 hours to degrade pre-existing DamPOIs before day 0. Dam activation was induced by 1 µM 4OH-TAM (Sigma, H6278) from day 8 ot day 10. Subsequently, cells were washed multiple times to thoroughly remove residual 4OH-TAM and cultured with 5 µM dTAG-13 to degrade Dam protein until day 17. For the Day-17 snapshot condition, 4-OHT was applied on days 16–17.

### Animals

Male and female C57BL/6JRJ mice (Janvier, France) were acclimated to the vivarium for at least 5 days before the experiments. Adult animals were group-housed (3–4 per cage) with enriched conditions in a 12 hr light/dark cycle (the light switches on at 6 AM) with a constant level of humidity and temperature (22 ± 1 °C). Food and water were provided ad libitum. Pregnant females were single-housed during the last 5 days of pregnancy. Pups were weaned at postnatal day 21. All the experimental procedures were conducted according to the Danish Animal Experiment Inspectorate.

### AAV injection

Mice were anesthetized using isoflurane (IsoFlo vet 100%, Zoetis), and they were kept on a heating pad for the duration of the procedure. Preoperative care included application of eye ointment for lubrification and subcutaneous injection with buprenorphine 0.3 mg/mL (Temgesic, 0.1 mg/kg) for analgesia. Standard surgical procedures were used to expose the skull and to identify the injection sites. All the coordinates are related to Bregma unless specified otherwise.

### Mice were bilaterally injected with the following mixture

AAV-9/2-CMV-promoter::rtTA-2A-CreERT2 **(titer:** 3.8 x 10^13^ vg/ml) mixed with *AAV-9/2- TRE-Tight2-promoter::LoxP-miRFP670nano-FKBP12^F36V^-Dam-Leo1* (titer: 6.1 x 10^12^ vg/ml) with *AAV-5/2-mCaMKIIα-promoter::mCherry* (titer: 6 × 10^12^ vg/mL) at the following ratio: 1:2:2. The injection volume was 0.25 µL per hemisphere at the following coordinates for primary visual cortex (V1, for bulk DamID-seq): AP: –4.0 mm; ML: ±2.9 mm; DV: −0.8/–1.0 mm from the skull. A different subset of mice was injected bilaterally in the medial prefrontal cortex (mPFC, for scDam&Tag) at the following coordinates: AP: +2.1 mm; ML: ±0.4 mm; DV: –1.5 mm from the skull with the same viral mix.

AAV-9/2-CMV-promoter::rtTA-2A-CreERT2 **(titer:** 3.8 x 10^13^ vg/ml) mixed with *AAV-9/2- TRE-Tight2-promoter::LoxP-miRFP670nano-FKBP12^F36V^-Dam-Laminb1* (titer: 1.4 x 10^13^ vg/ml) with *AAV-5/2-mCaMKIIα-promoter::mCherry* (titer: 6 × 10^12^ vg/mL) in sterile PBS at the following ratio: 1:2:2:9. Mice were injected bilaterally in V1 at the same coordinates and volume as described above.

AAV-9/2-CMV-promoter::rtTA-2A-CreERT2 **(titer:** 3.8 x 10^13^ vg/ml) mixed with *AAV-9/2- TRE-Tight-promoter::LoxP-FLAG_FKBP12^F36V^_EGFP* (titer: 2.0 x 10^13^ vg/ml) with *AAV- 5/2-mCaMKIIα-promoter::mCherry* (titer: 6 × 10^12^ vg/mL) at the following ratio: 1:1:1. Mice were injected bilaterally in V1 at the same coordinates and volume as described above.

Pups underwent a surgical procedure on postnatal day 1 or 2. They were anesthetized by placing them on an ice-cooled aluminium plate for 1-2 minutes until the pup was fully anesthetized. The right lateral ventricle was identified at 2/5 of the distance from the lambda suture to right eye as described^68^. By using a Hamilton precision syringe (662RNR; 600 series, 2.5 µL, needle size 22s ga) connected to a polyethylene ending with a 33 G injector, mice were injected with 2 µL of the following mixture:

AAV-9/2-CMV-promoter::rtTA-2A-CreERT2 **(titer:** 3.8 x 10^13^ vg/ml) mixed with *AAV-9/2- TRE-Tight2-promoter::LoxP-miRFP670nano-FKBP12^F36V^-Dam-Leo1* (titer: 6.1 x 10^12^ vg/ml) with *AAV-PHP.eB/2-shortCAG-promoter::-tdTomato* (titer: 1.4 × 10^13^ vg/mL) at the following ratio: 1:2:1 mixed with Evans Blue (Thermofisher, 195550050) at a final concentration of 0.05%. Evans Blue was used to ensure that the virus injection was delivered into the ventricles.

*AAV-9/2-CMV-promoter::rtTA-2A-CreERT2* **(titer:** 3.8 x 10^13^ vg/ml) mixed with *AAV-9/2- TRE-Tight2-promoter::LoxP-miRFP670nano-FKBP12^F36V^-Dam-Laminb1* (titer: 1.4 x 10^13^ vg/ml) at the following ratio: 1:2.5 mixed with Evans Blue at a final concentration of 0.05%.

### *In vivo* drug treatment

Dox (75 mg/kg, 450 uL; Sigma, D9891) was administered once by intraperitoneal injection in adult mice. To allow transgene expression in neonates, Dox was injected into the mother twice in a day (8 hours apart) as described^69^. 4OH-TAM (35 mg/kg; Sigma, H6278) was administered by intraperitoneal injection three times (once per day on alternating days). 4-Hydroxytamoxifen was dissolved as described^70^.

### Tissue collection and dissections

Tissues were collected from wild-type mice representing a random mix of males and females across different ages. Mice were first anesthetized with an overdose of isoflurane, followed by decapitation. Brains were submerged in ice-cold Phosphate-buffered saline. For the mice injected in V1, the area was identified under a fluorescent microscope. The left and right V1 areas, which were mCherry positive, were collected separately and snap frozen. Tissue was stored at -80 °C until further usage. For the mice injected in the ventricle at P-1/2, the cortical areas were separated from the subcortical areas. Only cortical areas were used for sequencing experiments. Cortical areas from the left and right hemispheres were collected separately, and cortical areas anterior to bregma were separated from the areas posterior to bregma.

### Smart-seq2

Total RNA was extracted using the RNeasy Mini Kit (Qiagen, 74104) with on-column DNase I treatment. RNA concentration was assessed using Nanodrop. cDNA synthesis and library preparation were performed using a modified Smart-seq2 protocol^71^. RNA (up to 1 ng) was mixed with 1 µL of 10 mM dNTPs (Thermofisher, 18427013) and 0.1 µL of 100 µM oligo-dT₃₀VN (synthesized by Sigma), heated at 72 °C for 3 min, chilled on ice, and then reverse transcribed with SuperScript II (Thermofisher, 18064022) in a 5.4 µL reaction containing 5X First-Strand Buffer, 100 mM DTT, 5 M betaine (Ampliqon, A351104), 1 M MgCl₂ (Thermofisher, 10418464), 100 µM template-switching oligo (TSO; synthesized by Qiagen), and RNase inhibitor (Promega, N2511). Reverse transcription was performed at 42 °C for 90 min, followed by 10 cycles of 50 °C for 2 min and 42 °C for 2 min, then 72 °C for 15 min. cDNA was pre-amplified by adding 15 µL of a preamplification mix containing KAPA HiFi HotStart ReadyMix (Roche Diagnostics, KK2602) and 100 µM ISPCR primer (synthesized by Sigma), using the following cycling conditions: 98 °C for 3 min; 12–15 cycles of 98 °C for 20 s, 67 °C for 15 s, and 72 °C for 6 min; and a final extension at 72 °C for 5 min. Amplified cDNA was purified with 1× Ampure XP Beads (Beckman, A63881) and eluted in 10 µL of 10 mM Tris-HCl (pH 7.6). For library preparation using in-house Tn5 transposase (12.5 µM, loaded with annealed adapters, Supplementary Table), 5 µL of 0.2 ng/µL purified cDNA was mixed with 15 µL tagmentation buffer (prepared using 5× TAPS-DMF buffer containing 50 mM TAPS pH 8.5 (Thermofisher, J63268.AE), 25 mM MgCl₂, and 50% Dimethylformamide (VWR, TCIAD0722)), incubated at 55 °C for 7 min, and the reaction was terminated with 5 µL of 0.2% SDS. After incubation at 25 °C for 7 min, 22.5 µL of library PCR mix containing Phusion polymerase (Thermofisher, F530S), dNTPs, and buffer was added along with 2.5 µL of IDT for Illumina DNA/RNA UD Indexes (Illumina, 20027213). PCR was performed with the following conditions: 72 °C for 3 min, 95 °C for 30 s, 10 cycles of 95 °C for 30 s, 55 °C for 30 s, and 72 °C for 30 s, followed by 72 °C for 5 min. Libraries were purified using Ampure XP Beads and eluted in 12 µL of 10 mM Tris-HCl (pH 7.6). Concentrations were determined using Qubit (Thermofisher), and the library fragment size was assessed using a Bioanalyzer (Agilent).

### DpnI-digestion-dependent DamID-seq

Genomic DNA was extracted from *in vitro* cultured cells and *in vivo* brain tissues by lysis in buffer containing 10 mM Tris-HCl, 100 mM NaCl, 10 mM EDTA, 0.5% SDS, and 400µg/mL Proteinase K at 56 °C for 1–3 hours. Following lysis, genomic DNA was purified by phenol:chloroform:isoamyl alcohol extraction, followed by ethanol precipitation. For DamID library preparation, up to 1 µg of genomic DNA was used. Purified genomic DNA was first digested with MboI (NEB, R0147L) restriction enzyme at 37 °C for 20 minutes, followed by heat inactivation. The DNA was dephosphorylated using Quick CIP (NEB, M0525L) to remove 5′-phosphates from DNA fragment ends. After heat inactivation of CIP, a second digestion was performed using DpnI (NEB, R0176L), followed by heat inactivation. To generate A-overhangs, the DpnI-digested DNA was treated with Klenow (exo–) (NEB, M0212) in the presence of dATP (NEB, N0440S). A-tailed DNA fragments were purified using AMPure XP Beads. Note that we did not use NEBNext Ultra II End Repair module. Adaptor ligation was performed using the NEBNext Ultra II Ligation Module (NEB, E7595), and library amplification was carried out using the NEBNext Ultra II Q5 Master Mix (NEB, M0544) and NEBNext Multiplex Oligos for Illumina (NEB, E6440), according to the manufacturer’s instructions. The library was subjected to size selection using a two-step AMPure XP Bead cleanup (0.6× followed by 1.0×) and assessed for quality using Qubit and Bioanalyzer. Dam-off (dTAG-13+) mESC shows a very low yield of library DNA without detectable peaks in Bioanalyzer, which accordingly resulted in very low read depth after high-throughput sequencing.

### Bulk Dam&Tag

Harvested cells were collected by centrifugation at 300 × g for 3 min at room temperature, resuspended in CELLBANKER1 (ZENOGEN PHARMA), and cryopreserved at –80 °C. Cryopreserved cells were thawed at 37 °C, immediately transferred to ice-cold 2% FBS in Pre-Wash Buffer (20 mM HEPES pH 7.5, 150 mM NaCl) and centrifuged at 300 × g for 3 min at 4 °C. Cells were then resuspended in 100 µL of ice-cold Antibody Buffer (Pre-Wash Buffer supplemented with 2 mM EDTA, 0.01% digitonin (Thermofisher, BN2006), 0.01% NP-40 (Thermofisher, 85124), 0.5 mM spermidine, 1% BSA (Sigma, A9647), and 1× protease inhibitor (Sigma, 11836170001)) with or without 1:500 m6A-Tracer-V5-EGFP (in-house preparation, 23 µM), and incubated for 1–2 h at 4 °C with rotation. After centrifugation (300g 3min 4 °C), cells were incubated with 100 µL of Antibody Buffer containing 1 µg anti-V5 or EGFP antibody (see Supplementary Table 3 and Extended Data Fig. 4a) overnight at 4 °C. After centrifugation, cells were incubated in 100 µL of Antibody Buffer with 1 µg secondary antibody (see Supplementary Table 3) for 1 h at room temperature with rotation, followed by two washes in 300 mM NaCl Buffer (Pre-Wash Buffer supplemented as above, adjusted to 300 mM NaCl, and EDTA removed). Cells were then incubated for 1 h at room temperature with 50 µL of 300 mM NaCl Buffer containing 1:25 Protein A-Tn5 (in-house loaded with ME adapters at 12.5 µM), followed by two additional washes with the same buffer. Tagmentation was performed in 50 µL of Tagmentation Buffer (300 mM NaCl Buffer with 10 mM MgCl₂ (Thermofisher, AM9530G)) at 37 °C for 1 h with shaking at 300 rpm. DNA was purified using the MinElute PCR Purification Kit (Qiagen, 28004) and eluted in 10 µL of EB buffer. Libraries were prepared using Q5 High-Fidelity 2X Master Mix (NEB, M0492) and IDT for Illumina DNA/RNA UD Indexes Set A (Illumina, 20091654). Each 25 µL PCR reaction contained 2.5 µL indexed primers and 12.5 µL Q5 Master Mix. Thermal cycling conditions were: 72 °C for 5 min, 98 °C for 30 sec, followed by 10–12 cycles of 98 °C for 10 sec, 63 °C for 30 sec, and 72 °C for 1 min, with a final extension at 72 °C for 1 min. Half of the PCR product was purified with 0.6× AMPure XP Beads and eluted in 20 µL of 10 mM Tris-HCl (pH 7.6). Library yield and size were assessed using Qubit and Bioanalyzer, and if the concentration was below 2 ng/µL, additional PCR cycles were performed on the remaining PCR products. V5- EGFP-m6A-Tracer-minus protocol resulted in a very low yield of library DNA (see Extended Data Fig. 4b) without detectable peaks in Bioanalyzer, which accordingly resulted in very low read depth after high-throughput sequencing.

### scDam&Tag

Flash-frozen tissue samples (mPFC from Dam-labeled mice) were first homogenized (VWR, 432-1270) to isolate nuclei. Prior to homogenization, a Dounce homogenizer was chilled on ice, and forceps for tissue handling were cooled on dry ice. The frozen tissue was rapidly transferred into a pre-chilled 1 mL homogenizer, and 500 µL of ice-cold Antibody Buffer was immediately added. The tissue was homogenized using 5 strokes with the loose pestle, followed by 10–15 strokes with the tight pestle, keeping all steps on ice. The homogenate was filtered through a 30 µm strainer (Miltenyi Biotec, 130041407) into a 1.5 mL tube, centrifuged at 300 g for 3 min at 4 °C (e.g., swing rotor of Eppendorf 5810R), and the supernatant was removed. The pellet was resuspended in 500 µL of Antibody Buffer containing m6A-Tracer-V5-EGFP (23 µM, 1:500 dilution) and incubated for 1 hour at 4 °C with rotation. After centrifugation at 300 g for 3 min at 4 °C and removal of the supernatant, the nuclei were incubated overnight at 4 °C with mouse anti-V5 antibody (mouse monoclonal, Thermorisher, R960-25, 1 µg/µL, 1:100 dilution) diluted in 500 µL of Antibody Buffer. This was followed by a 1-hour incubation with anti-mouse secondary antibody (Abcam, ab46540; 1:100 dilution) in 500 µL of Antibody Buffer at room temperature with rotation. After washing, nuclei were incubated with in-house protein A-Tn5 transposase (12.5 µM, 1:10 dilution; ME-A adaptor with sample-barcode (Supplementary Table 1)) that had been pre-loaded with a unique index adaptor specific to each sample. This was performed in 200 µL of ice-cold 300 mM NaCl Buffer for 1 hour at room temperature with rotation. Nuclei were washed twice with 500 µL of 300 mM NaCl Buffer, then subjected to tagmentation in 200 µL of MgCl₂ + Tagmentation Buffer at 37 °C for 1 hour with shaking at 300 rpm. The reaction was stopped with ice-cold DNB Stop Buffer (1× DNB (20x Diluted Nuclei Buffer, 10x Genomics), 25 mM EDTA, and 1% BSA). The nuclei were pelleted by centrifugation at 300 g for 2 min at 4 °C, washed once with ice-cold 1% BSA DNB, and stained with 7-AAD (1:1000 dilution) in 1 mL of ice-cold 1% BSA DNB. Nuclei were filtered through a 50 µm strainer and immediately sorted by FACS to isolate m6A-GFP⁺/7-AAD⁺ populations into ice-cold 2% BSA DNB. Equal numbers of nuclei from two biological samples were pooled, centrifuged at 500 g for 5 min at 4 °C, and resuspended in 1% BSA DNB to a final concentration of ∼4,000 nuclei/µL. A total of 32,000 nuclei (i.e., 16,000 per sample) were used for the 10x Genomics Chromium Next GEM Single Cell ATAC v2 (1000406) workflow, including GEM generation, barcoding, and linear amplification, following nanoCT protocol^4,65^. Specifically, 8 µL of nuclei suspension was mixed with 7 µL of ATAC Buffer (10x Genomics) and transferred to 60 µL of Barcoding Master Mix (56.5 µL Barcoding Reagent B, 1.5 µL Reducing Agent B, and 2 µL of Phusion High-Fidelity DNA Polymerase (Thermofisher, F530S) in place of Barcoding Enzyme to increase the complexity of library). GEM generation was performed using the 10x Chromium Controller, followed by linear amplification PCR with the following conditions: 72 °C for 5 min, 98 °C for 30 sec, then 12 cycles of 98 °C for 10 sec, 59 °C for 30 sec, and 72 °C for 1 min, with a final extension at 72 °C for 1 min. The post-GEM PCR product was followed by phase separation using Recovery Agent and bead-based cleanup (MyOne SILANE beads + Cleanup Buffer + Reducing Agent). Eluted DNA in 40.5 µL of Elution solution (98 µL of EB Buffer (QIAGEN), 1 µL 10% Tween 20, and 1µL Reducing Agent B) was further purified using AMPure XP Beads and eluted in 40 µL EB buffer. DNA concentration was quantified with a Qubit fluorometer to adjust the amount of Tn5 used in the subsequent step. DNA was subjected to a second tagmentation using in-house Tn5 loaded with ME-B-only adaptors (12.5 µM) in 40 µL of 2× TrisTD Buffer (20 mM Tris-HCl, pH 7.6, 10 mM MgCl₂, 20% DMF) at 37 °C for 30 min. After cleanup with 128 µL of AMPure XP Beads, DNA was eluted in 40 µL of EB Buffer. For final library amplification, a pre-incubation step was performed in which 50 µL of Amp Mix and 7.5 µL of SI-PCR Primer B were heated at 98 °C for 1.5 min. The mixture was then cooled to below 72 °C before 40 µL of DNA sample and 2.5 µL of Single Index N Set A (10x Genomics, 1000212) were added, and PCR amplification was performed with the following thermal profile: 72 °C for 1 min, 98 °C for 45 sec, then 13 cycles of 98 °C for 20 sec, 67 °C for 30 sec, and 72 °C for 30 sec, followed by a final extension at 72 °C for 1 min. Half of the final library was subjected to size selection using a two-step AMPure XP bead cleanup (0.4× followed by 1.1×) and assessed for quality using Qubit and Bioanalyzer.

### scHisTrac-seq using nanobody-Tn5

Six experimental groups were prepared: Day-2 and Day-7 differentiated neurons (snapshots), and Day-2 -> Day-7 HisTrac, each with two biological replicates. Cryopreserved cells stored in CELLBANKER1 were thawed in a 37 °C water bath, immediately diluted with 500 µL of 2% FBS/Pre-Wash buffer, filtered through a 30 µm cell strainer, and centrifuged at 300 × g, 4 °C, for 3 min. Cell pellets were resuspended in 200 µL of Antibody Buffer per sample containing m6A-Tracer-V5-EGFP (23 µM stock, 1:500 dilution) and incubated at 4 °C for 1 h with rotation. After centrifugation and removal of the supernatant, cells were incubated in 200 µL of Antibody Buffer containing anti-V5 (mouse monoclonal, Thermorisher, R960-25, 1 µg/µL, 1:100 dilution) and anti-H3K27ac (rabbit polyclonal, Abcam ab4729, 0.4 µg/µL, 1:40 dilution) antibodies overnight at 4 °C with rotation. The next day, samples were centrifuged, and the supernatant was removed. Nuclei were then incubated for 1 h at room temperature in 200 µL of 300 mM NaCl buffer containing 10 µL each of in-house prepared Mnb-Tn5 and Rnb-Tn5 (12.5 µM), which are nanobody-Tn5 transposase complexes targeting mouse or rabbit IgG, respectively. Each nanobody-Tn5 was pre-loaded ME-A adapter with 12 unique sample/modality-specific barcodes (Supplementary Table 1). After removal of supernatant by centrifugation, samples were resuspended in 200 µL of 300 mM NaCl buffer and incubated with rotation at 4 °C for 20 min. Following centrifugation, a second wash was performed with fresh 300 mM NaCl buffer and immediate centrifugation. Tagmentation was initiated by adding 50 µL of MgCl₂+ Tagmentation Buffer and incubating at 37 °C for 1 h with shaking at 300 rpm. The reaction was stopped with 50 µL of 1% BSA-containing DNB Stop Buffer (1× DNB, 1% BSA, and 25 mM EDTA), followed by sequential washes in 1% BSA DNB.

After centrifugation, nuclei were resuspended in 400 µL of ice-cold 1% BSA/DNB containing 7-AAD (1:1000 dilution), filtered through a 50 µm strainer, and sorted by FACS. 7-AAD⁺/m6A-GFP⁺ nuclei were collected into 2% BSA/DNB. Nuclei from different samples were pooled and centrifuged (500 × g, 4 °C, 5 min), and the supernatant was removed. Nuclei were resuspended in 1% BSA/DNB. A maximum of 96,000 nuclei (i.e., 16,000 per sample) were used for the 10x Genomics Chromium Next GEM Single Cell ATAC v2 workflow, including GEM generation, barcoding, and linear amplification, following nanoCT protocol^4,65^. Specifically, 8 µL of nuclei suspension was mixed with 7 µL of ATAC Buffer (10x Genomics) and transferred into 60 µL of Barcoding Master Mix, consisting of 56.5 µL Barcoding Reagent B, 1.5 µL Reducing Agent B, and 2 µL of Phusion High-Fidelity DNA Polymerase (in place of the standard Barcoding Enzyme). GEM generation was performed using the 10x Chromium Controller, followed by linear amplification PCR under the following thermal conditions: 72 °C for 5 min, 98 °C for 30 sec, then 12 cycles of 98 °C for 10 sec, 59 °C for 30 sec, and 72 °C for 1 min, with a final extension at 72 °C for 1 min. Post-GEM reactions were subjected to phase separation using Recovery Agent and cleaned up with MyOne SILANE beads in the presence of Cleanup Buffer and Reducing Agent. Eluted DNA in 40.5 µL of Elution Solution (98 µL of EB Buffer (QIAGEN), 1 µL 10% Tween 20, and 1µL Reducing Agent B) was further purified using AMPure XP beads and eluted in 41.5 µL of EB Buffer. DNA concentration was quantified with a Qubit fluorometer to adjust the amount of Tn5 used in the subsequent step. The secondary tagmentation was performed using in-house Tn5 loaded with ME-B-only adapters (12.5 µM) in 40 µL of 2× TrisTD Buffer (20 mM Tris-HCl, pH 7.6, 10 mM MgCl₂, 20% DMF) at 37 °C for 30 min. Following cleanup with 128 µL of AMPure XP beads, DNA was eluted in 40 µL of EB Buffer. Final libraries were PCR-amplified using 50 µL of NEB Q5 High-Fidelity 2× Master Mix, 7.5 µL of SI-PCR primer, and 2.5 µL of Single Index N Set A. PCR conditions were: 72 °C for 5 min, 98 °C for 30 sec, then 13 cycles of 98 °C for 20 sec, 67 °C for 30 sec, and 72 °C for 1 min, followed by a final extension at 72 °C for 1 min. Half of the final amplified library was size-selected using a two-step AMPure XP bead cleanup (0.4× followed by 1.1×), and quality was assessed using a Qubit fluorometer and Bioanalyzer.

### Sequencing

Smart-seq2 and bulk DamID libraries were sequenced for paired-end with NovaSeq X plus (Illumina) and DNBSEQ (BGI). scDam&Tag and scHisTrac with sample/modality- barcoded ME-A adapters were sequenced for paired-end with NovaSeq X plus (R1-i7-i5- R2 = 40-8-41-40) using custom sequencing primers (i.e., Seq_primer_R1 and Seq_primer_I2; Supplementary Table 3), which were designed following previous studies^4,65^.

### Fluorescence-activated cell sorting (FACS)

After tagmentation of nuclei in scDam&Tag and scHisTrac-seq protocols, in which Dam-POI-overexpressing nuclei were stained with V5-EGFP-m6A-Tracer, entire nuclei were further labeled with 7-Aminoactinomycin D (7-AAD; Thermofisher, 10748784). We used a 6-laser Bigfoot cell sorter (Thermofisher) for sorting single EGFP-expressing nuclei. 7- AAD signal was used to gate intact nuclei, a forward scatter area versus height followed by a side scatter area versus height plot was used to sort singlets, and EGFP was used to define Dam-POI-overexpressing nuclei for sorting (gates are shown in Extended Data Figs. 5 and 6).

### Staining

The mice were anesthetized with isoflurane and euthanized by cervical dislocation. The brains were harvested and stored for 24 hr in 10% formalin at room temperature on a shaker. Then, the brains were sliced into 120-µm-thick slices in PBS on a Leica Vibratome (VT1000S).

Slices were permeabilized with PBS-Triton X-100 0.5%, 10% of normal goat serum (NGS), and blocked in 10% bovine goat serum albumin (BSA) for 90 min at room temperature. Subsequently, the slices were incubated with anti-GFP (Thermofisher, CAB4211, 1:1000) in PBS-Triton X-100 0.3%, 1% NGS, and 5 % BSA, and the incubation lasted for 72 hr at 4°C. At the end of the incubation time, the slices were washed three times in PBS. The slices were incubated in Alexa Fluor 488 goat anti-rabbit (Thermofisher, A-11008, 1:1000) in PBS-Triton X 0.3%, 1% NGS, and 5% BSA for 24 hr at 4°C. Nuclear staining was performed by using 1:1000 of DAPI (Sigma, D9542) for 30 min at room temperature. Brain slices were mounted on poly-lysine glass slides with coverslips using Fluoromount G (Southern Biotech).

### Imaging and cell counting

Imaging was performed by using a virtual slide scanner (Olympus VS120). Tile images were taken by the whole brain slides by using ×10 (UPLSAPO 2 ×10/0,40) objective). The emission wavelength for Alexa 488 was 518 nm with 3.2 ms of exposure time. The emission wavelength for DAPI was 455 nm with 3.2 ms of exposure time. GFP-positive cells were counted manually by using ImageJ. The experimenter, blind to the treatment, defined a region of interest and performed the cell counting. The GFP-positive cells were quantified in V1 bilaterally at AP: –4.0 mm from bregma, the region with the highest density of GFP-positive cells in the control group. The area dimension, used for normalization, was calculated by using the built-in function in ImageJ.

### Bulk sequencing data processing

#### Smart-seq2 RNA-seq data processing

Raw paired-end reads were adapter-trimmed using *cutadapt* (v3.5) with the following parameters: adapter sequence CTGTCTCTTATACACA on both 5′ and 3′ ends. Trimmed R1/R2 pairs were then aligned to the mouse genome (mm10) using *STAR*^72^ (v2.7) in two-pass mode with ENCODE-recommended settings. Aligned BAM files were indexed with *samtools*^73^ (v1.6).

#### Bulk DamID-seq data processing

Raw paired-end reads were first filtered to enrich for genuine DpnI-generated fragment ends by retaining only read pairs whose R1 and R2 sequences begin with a 5′ “TC” dinucleotide (the overhang produced by DpnI digestion). This filtering was performed with *cutadapt* (v3.5) using anchored 5′ adapters (*-g ^TC -G ^TC*), discarding any read pairs lacking the TC motif (*--discard-untrimmed*), and retaining only fragments ≥10 nt post-filtering (*-m 10*). Filtered reads were then trimmed to remove the standard Illumina adapter sequence. Trimmed, TC-filtered read pairs were aligned to the mm10 reference genome using *Bowtie2*^74^ (v2.5.1). Sam output was piped through *samtools* (v1.6) to produce coordinate-sorted BAM files.

#### Bulk Dam&Tag and ATAC-seq data processing

Raw paired-end reads were adapter-trimmed using *cutadapt* (v3.5) with the following parameters: adapter sequence CTGTCTCTTATACACA on both 5′ and 3′ ends. Trimmed read pairs were aligned to the mm10 reference genome using *Bowtie2* (v2.5.1). Sam output was piped through *samtools* (v1.6) to produce coordinate-sorted BAM files.

#### Public ChIP-seq data processing

Public datasets of ChIP-seq against H3K27ac, H3K4me3, H3K27me3, and Laminb1 in mESC were first processed by *cutadapt* (v3.5) to remove adapter sequence and aligned to the mm10 reference genome using *Bowtie2* (v2.5.1). Sam output was piped through *samtools* (v1.6) to produce coordinate-sorted BAM files.

### Enhancer identification

Trimmed, aligned ATAC-seq BAM files from mESC, Day-2 neurons, and Day-7 neurons were processed with *MACS2*^75^ (v2.2.7.1) in paired-end mode (*-f BAMPE*) to call open-chromatin peaks. All biological replicates were supplied together, using the mouse genome size parameter (*-g mm*). Peak summits were imported into R as a GRanges object, and each summit was extended ±500 bp around its midpoint to define peak regions. To distinguish enhancers from promoters, ATAC-peaks were compared against annotated TSS regions (TSS ±1 kb) from TxDb.Mmusculus.UCSC.mm10.knownGene. TSS non-overlapping ATAC-peaks were designated putative enhancers.

### BigWig preparation and genome browser views

BAM files were imported into a *QuasR* (1.44.0) in *R* (4.4.0) via *qAlign*. Genome-wide coverage was exported to BigWig format using *qExportWig* at 100 bp to 1 Kbp resolution. Coverage values were scaled to counts per million mapped reads to normalize for library size. We obtained Free-Dam-normalized coverage tracks with the *bigwigCompare* function of *deepTools*^76^ (v.3.5.5). We compared each Dam-POI BigWig to its matched Free-Dam BigWig. We computed the log₂ ratio of normalized coverage (*--operation log2*), adding an optimized pseudocount to avoid division by zero (*--pseudocount*), and output the result as a BigWig (*--outFileFormat bigwig*). We used the IGV genome browser (v2.16.0) for visualization.

### Smart-seq2 gene expression quantification and statistics

Smart-seq2 data was quantified at the gene level using the *QuasR*^77^ in *R*: reads were quantified over exons annotated in the TxDb.Mmusculus.UCSC.mm10.knownGene transcript database and summarized to genes using *qCount*. Subsequently, RPKM (reads per kilobase of exon per million mapped reads) was quantified. For differential expression, raw counts were loaded into *edgeR*^78^ (v4.2.2). Genes with very low expression were removed using *filterByExpr*, and library-size normalization factors were computed by the TMM method (*calcNormFactors*). Common, trended, and tagwise dispersions were estimated sequentially (*estimateGLMCommonDisp*, *estimateGLMTrendedDisp*, *estimateGLMTagwiseDisp*). A negative-binomial generalized linear model was then fit (*glmFit*). Differential expression between Dam–POI-on and off conditions was tested with the likelihood-ratio test (*glmLRT*), and genes with a Benjamini–Hochberg FDR < 0.01 and absolute log₂–fold change > 0 (i.e., no fold-change cut-off) were considered significant.

### Bulk ATAC-seq and ChIP-seq quantification

Aligned reads were quantified genome-wide using the *QuasR* package. m6A signal was quantified at the levels of promoters (TSS ± 3 Kbp), enhancers, or genomic bins (100 Kbp for Laminb1, 10 Kbp for other signatures) using *qCount*. Subsequently, RPKM was quantified.

### Bulk DamID-seq and Dam&Tag quantification and statistics

Aligned reads were quantified genome-wide using the *QuasR* package in *R*. m6A signal was quantified at the levels of genes (TSS-TES), promoters (TSS ± 3 Kbp), enhancers, or genomic bins (100 Kbp for Dam-Laminb1, 10 Kbp for other Dam-POIs) using *qCount*. We retained only genomic features containing at least one GATC motif to focus on the m6A signal, and counts per million mapped reads (CPM) were computed. Dam-POIs other than Dam-RNAPII and Dam-Leo1 were normalized to the Free-Dam to control locus-specific variation in DamID: log₂–ratio of Dam-POI signal over Free-Dam signals was computed. Regarding the genomic feature-size normalization, because the number of GATC sites varies across genomic features, and thus the opportunity for m6A modification differs, we normalized counts per million (CPM) by the local GATC density, rather than the length of genomic features.

Regarding differential testing between Day-2 and Day-7 samples for Dam-Leo1, Dam- RNAPII, and *in vivo* Dam-Laminb, raw m6A counts for the two biological replicates per stage were imported into *edgeR*. Genomic features with low coverage were filtered out, and the remaining data were normalized for library size using the TMM method (*calcNormFactors*). Common, trended, and tagwise dispersions were estimated sequentially (*estimateGLMCommonDisp*, *estimateGLMTrendedDisp*, *estimateGLMTagwiseDisp*). A negative-binomial generalized linear model was then fit (*glmFit*), and the contrast was tested by a likelihood-ratio test (*glmLRT*). Benjamini– Hochberg FDR < 0.01 and absolute log₂–fold change > 1 were used for selection. Regarding the clustering, pairwise sample distances of log₂-CPM values were computed using Euclidean distance, and sample correlations were calculated using Pearson’s method, and a corresponding distance metric (1 – correlation) was clustered with complete linkage. They were visualized by the *pheatmap* package (v1.0.13).

Regarding differential testing between Day-2 and Day-7 samples for Dam-Laminb1, Dam- Taf3, Dam-mH3K27me3, and Dam-Biv, GATC- and Free-Dam–normalized DamID counts were used. Features with minimal signal (log₂ signal ≤ 0.3 in all samples) were removed. A linear model was fit to the data via *lmFit* (*limma*^79^ v3.60.0), and the pairwise contrast was defined with *makeContrasts* and applied using *contrasts.fit*. Empirical Bayes moderation (*eBayes*) improved variance estimates. Differentially enriched features were identified from the resulting empirical Bayes–moderated *t*-statistics. Benjamini–Hochberg FDR < 0.01 and absolute log₂–fold change > 1 were used for selection. Regarding the clustering, pairwise sample distances were computed using Euclidean distance, and sample correlations were calculated using Pearson’s method, and a corresponding distance metric (1 – correlation) was clustered with complete linkage. They were visualized by the *pheatmap* package.

To assess the correlation between mRNA levels and Dam-POIs (i.e., Leo1, RNAPII, FreeDam) -mediated m6A deposition, we calculated the average RPKM across three Smart-seq2 mESC replicates and the average GATC-normalized Dam-POI signals across two biological replicates. For each gene, means were computed and log₂- transformed with a pseudocount, and then filtered out genes with low signals. Pearson’s correlations were calculated.

### Aggregate plots and statistics

We first prepared four BED files of genomic features (i.e., genes, enhancers, 100 Kb genomic bins, and promoters) based on their levels of mRNA, ATAC-seq, ChIP-seq H3K27ac, ChIP-seq H3K4me3, ChIP-seq H3K27me3, and ChIP-seq Laminb1 in mESCs (top 25%, 50%, 75% and 100% by RPKM). For each mESC Dam-POI BigWig track of interest, we computed a matrix of binned coverage using *deepTools*’ *computeMatrix* scale-regions (only for genes) or *reference-point* (other genomic features). Zero-coverage features were skipped. Aggregate profiles were plotted with *plotProfile*, using the “*se*” plot type to display the mean ± standard error across genes.

### Demultiplexing of single-cell Dam&Tag and HisTrac-seq

We used the *nanoscope* pipeline, designed for the analysis of nanoCT^4,65^. Raw sequencing reads were first demultiplexed by sample/modality-specific barcodes into separate FASTQ files. These FASTQ files were then processed with *Cell Ranger ATAC* (v2.0.0) to perform alignment to the mm10 reference genome, basic quality control, and preparation of fragments files (fragments.tsv.gz). Next, broad-peak calling was executed on aggregated pseudo-bulk fragments for each modality using *MACS2*. Finally, *nanoscope* collated all output into a unified metadata table, annotating each cell with its total fragments, unique fragment counts per cell, and FRiP for downstream analysis. In our implementation of the *nanoscope* demultiplexing script (debarcode.py), we replaced the original antibody-barcode “follow” pattern *default="GCGTGGAGACGCTGCCGACGA"* with *"GACGCTGCCGACGA"*. This change was necessary because our sample/modality-barcoded ME-A adapters are designed based on NTT-seq^49^, deviated from the nanoCT^4,65^, so the hard-coded 20-mer no longer matches the region immediately downstream of the antibody barcode (Supplementary Table 3).

### Single-cell Dam&Tag and HisTrac-seq analysis with *nanoscope*, *Signac,* and *Seurat*

We adopted the Nanoscope “Analysis using peaks” workflow using *Seurat*^80^ (v5.3.0) and *Signac*^81^ (v1.14.0) (https://github.com/bartosovic-lab/nanoscope) for downstream single-cell chromatin profiling. For each modality, broad peaks called by *MACS2* were imported as GRanges. We then merged replicates’ peak sets via *reduce()* to define a unified, nonredundant peak atlas per modality, as recommended by *nanoscope*. Regarding metadata-driven cell filtering, rather than using *nanoscope*’s internal cell-calling, we applied 10x-derived QC metrics from the *CellRanger* output. We built Fragment objects from fragments.tsv.gz and extracted counts over the consolidated peak sets using *Signac*’s *FeatureMatrix()*, and we created individual *Seurat* objects per sample/modality, storing peak counts and metadata. We carried out an additional filtering by retaining cells passing the 1st–99th percentile thresholds for unique fragment counts per cell and > 1st percentile for FRiP. We further restricted our analysis to cells that passed QC in both Dam and H3K27ac modalities for each biological replicate. Replicates of the same modality were merged using *merge()*, preserving sample identity via cell-ID prefixes. The TF–IDF normalization (*RunTFIDF*) was performed to weigh peaks by their rarity across cells. Top features were selected with *FindTopFeatures*, followed by a latent semantic indexing step (*RunSVD*) to reduce dimensionality. Nearest-neighbor graphs in the LSI space were constructed with *FindNeighbors*, and Louvain clustering was applied (*FindClusters*) at modality-specific resolutions. UMAP embeddings (*RunUMAP*) were computed on selected LSI dimensions, yielding two modality-specific maps. 1^st^ dimension of the LSI space was not used. The number of LSI dimensions and resolution used for clustering and UMAP were optimized to observe two subpopulations (i.e., id-T and id-U). Minor clusters other than them were removed from the downstream analysis. Cluster identities from the K27ac assay were annotated and then transferred to the Dam modality (and vice versa) via metadata merging, allowing direct comparison of cluster assignments across modalities. To approximate transcriptomic profiles from chromatin accessibility, we used *Signac*’s *GeneActivity()* to count peak signals within 2 kb upstream and downstream of gene bodies. The resulting matrices were added as new *Seurat* assays (i.e., “RNA.K27ac” and “RNA.Dam”), log-normalized with a scale factor equal to the median library size. Gene expression, module scores, and peak signals were visualized by *Vlnplot* and *FeaturePlot*.

### Pseudobulk clustering of modalities using LSI medoids

To compare sample-level relationships in the single-cell datasets, we computed pseudobulk medoids in PC (i.e., LSI) space for each modality and replicate. Specifically, for each sample (rep1 and rep2), we identified the single cell (LSI point) whose Euclidean distance to all other cells in that sample was minimal, using *dist()* (R *stats* package (v4.4.0)) and selecting the row with the smallest total distance. These medoids serve as robust representatives of each replicate’s chromatin profile. We tested several variations of LSI space (always without 1^st^ dimension), and confirmed that the trend is conserved. We then calculated pairwise Euclidean distances between medoids and clustered them with Ward’s linkage to reveal sample groupings, and visualized the results alongside a Pearson’s correlation heatmap of the medoid vectors.

### Analysis of cell identity transition and statistics

We used *plotConnectModal* function of *nanoscope* (in functions_scCT2.R) to visualize cell identity transitions from Dam modality to H3K27ac modality on UMAP by connecting single-cells with the same cell barcodes with lines across different modalities. Colore- code was based on Dam-modality. In addition, we constructed alluvial plots using *ggalluvial* (0.12.5) to highlight both conserved assignments and disruptive “identity jumps” between modalities. To test if identity mismatch increases in stage-mismatch HisTrac (Day2-Day7) as compared to snapshots (Day2-Day2 and Day7-Day7), for each replicate, we computed the percentage of cells whose Dam vs. H3K27ac cluster labels disagree.

Subsequently, we performed Welch’s one-sided *t*-test, *t.test(trt, ctrl, alternative="greater", var.equal=FALSE)* (R *stats* package (v4.4.0)) to test whether the mean mismatch rate in the retrospective (HisTrac) group is significantly higher than in the stage-matched snapshot group. Dam-Leo1, t = 9.30, df = 3.01, *P*-value = 0.0013, Free-Dam, t = 5.431, df = 2.05, *P*-value = 0.0152; Dam-Biv, t = 8.125, df = 3.16, *P*-value = 0.0016.

### Opposing medoid distance (OMD) analysis and statistics

All cells from the Day-2, Day-7, and HisTrac samples were merged into a single Seurat object sharing a common LSI embedding, ensuring that every cell’s coordinates are directly comparable. For each modality, we identified two major clusters (id-T and id-U) based on their respective Dam- and K27ac-derived annotations (see above). Within each cluster, we computed the medoid score in LSI space, namely the actual cell minimizing the sum of Euclidean distances to all other cells in the cluster. Next, for every cell, we calculated its Euclidean distance to both cluster medoids for both Dam and K27ac embeddings. This yields four distance metrics per cell. We then defined the OMD for each cell as its distance to the opposite cluster’s medoid (e.g., a Dam-cluster-T cell’s dist_contra_Dam is its distance to the Dam-cluster-U medoid). Cells were labeled “homo” if their Dam and K27ac cluster IDs agreed, or “hetero” if they differed. We compared the distributions of OMDs (on a log₂ scale) between homo and hetero cells for each modality using boxplots. In addition, we computed the median OMD among homo cells (*m*), then counted the number *k* of hetero cells whose distance fell below *m* out of the total hetero count *N*. We expressed this as a percentage (*k/N × 100%)* and derived exact 95% Clopper–Pearson confidence intervals using the *binom.test(k, N)* (*stats*). We tested several variations of LSI space (always without 1^st^ dimension), and confirmed that the trend is conserved.

### Differential gene and peak analysis and statistics

Using H3K27ac modality of Dam-Leo1 single-cell samples, we screened stage-specific marker genes between Day 7 and Day 2, setting the assay to the “RNA” (RNA.K27ac), and invoked Seurat’s *FindMarkers* (default two-sided Wilcoxon rank-sum test, *test.use = "wilcox", logfc.threshold = 0.25, min.pct = 0.10*). Genes with adjusted *P*-value < 0.01 were retained as Day-7_up or Day-7_down markers (Supplementary Table 2). Next, to screen identity-specific programs, we repeated the same *FindMarkers* workflow comparing cells annotated as id-U versus id-T within the combined Day-2/Day-7 set. Genes with adjusted *P*-value < 0.05 were retained as id-U_up and id-T_down markers (Supplementary Table 2). We computed cell-level gene activity for each of these four gene sets using *Seurat*’s *AddModuleScore*, yielding Dev_UPscore, Dev_DOWNscore, U_UPscore, and U_DOWNscore; we then defined a combined developmental gene module score (Dev_UPscore – Dev_DOWNscore) and an identity gene module score (U_UPscore – U_DOWNscore) for every cell.

To identify genes whose Dam-mediated historical signal associates specifically with disruptive identity jumps (i.e., cells whose Day2-Dam- and Day7-H3K27ac-derived cluster labels disagree) within the retrospective HisTrac sample, we again used Seurat’s *FindMarkers* on the Dam–Leo1 “RNA” assay. First, we labelled each cell by its identity transition patterns (i.e., T->T, T->U, U->T, U->U). We then ran two two-group Wilcoxon rank-sum tests (via *FindMarkers(test.use="wilcox", logfc.threshold=0.25, min.pct=0.10)*) comparing each jump cell against control cell (T->U vs. T->T and U->T vs. U->U). Genes with adjusted *P*-value < 0.05 were taken as jump-associated gene markers (Supplementary Table 2). We computed cell-level gene activity for each of these two gene sets using *Seurat*’s *AddModuleScore*. To analyze the enhancers associated with identity jumps, we followed the same approach but using the “peaks” assay rather than gene-activity counts. We run *Seurat*’s *FindMarkers(test.use="wilcox", logfc.threshold=0.25, min.pct=0.10).* Peaks with adjusted *P*-value < 0.05 were taken as jump-associated peaks (Supplementary Table 2). All *P*-values were Bonferroni-corrected as implemented by Seurat.

### Gene Ontology (GO) analysis

For each DEG list (e.g., Day-7_up, Day-7_down, identity-specific and jump-associated sets), we performed GO enrichment using the *R* package *clusterProfiler* (v4.12.6) with the *org.Mm.eg.db annotation*. Gene symbols were passed to *enrichGO(ont="BP", pAdjustMethod="BH", pvalueCutoff=0.05, qvalueCutoff=0.05, minGSSize=5, maxGSSize=800, readable=TRUE)*. Enriched terms were visualized with *dotplot()*, showing the top 20 categories by adjusted *P*-value.

### Transcription factor motif analysis

For jump-associated peaks, we performed transcription factor motif enrichment using *Signac*’s *FindMotifs* function against the *JASPAR2024* vertebrate database^82^. We scanned the mm10 genome for motif occurrences with motifmatchr and *TFBSTools*, computed background frequencies from all detected peaks, and assessed enrichment by a two-sided Fisher’s exact test (*Signac* default). The top 25 enriched motifs (ranked by Benjamini–Hochberg–adjusted *P*-value) were visualized via *MotifPlot*, overlaying motif logo densities on the aggregated peak accessibility.

## Data availability

Sequencing raw data and processed data used for this study are deposited in ArrayExpress and will be released to the public without restrictions. mRNA-seq (Smart-seq2), E-MTAB-15338; bulk Dam, E-MTAB-15336; scDam, E-MTAB-15341. Public ChIP-seq data for mESC are from GSE23943 (H3K4me3, H3K27me3), GSE24164 (H3K27ac), and GSE97878 (Laminb1).

## Code availability

All codes for the processing of bulk and single-cell Dam will be deposited on GitHub without restrictions by publication.

## Acknowledgements

We thank the members of the Kitazawa laboratory, especially I. Faress, K. Itoh, S.W. Andersen, and M. Vestergaard for their kind support of experiments and analysis. We thank T. Kim (Aarhus University) and C. Ruescher (University of Geneva) for critical reading of the manuscript. We thank M. Kjærgaard’s lab (Aarhus University) for the protein purification protocol. We thank J. Kind (Hubrecht Institute) for advice on DamID. We thank N. Yachie (University of British Columbia) for advice on OMD. We thank G. Castero-Branco (Karolinska Institute) and M. Bartosovic for the discussion regarding the adaptation of the nanoCT protocol. We acknowledge the Danish Single-Cell Examination Platform (CellX), especially L. Lin (Aarhus University), for providing access to 10x Controller. CellX was established with support from the Danish Research Agency through the Danish national research infrastructure program (5229-0009B). We appreciate that cell sorting was performed at the FACS Core Facility (Aarhus University). This study was supported by Lundbeckfonden (grant no. DANDRITE-R361-2020-2654, R436-2023-1201, and R436-2023-1201), European Research Council ERC-StG MemoPlasticGenomics (grant no. 101039734), Novo Nordisk Foundation (grant no. 0096135 and 0088740), Aarhus Universitets Forskningsfond (AUFF) (grant no. AUFF-E-2024-9-40), and Japan Science and Technology Agency (JST) PRESTO (grant no. JPMJPR24N6).

## Author contributions

T.K. and Y.K.K. designed the study; Y.K.K. performed most of the *in vitro* experiments; V.K. performed most of the *in vivo* experiments; Y.K.K., V.K., and T.K. analyzed data; All authors wrote the manuscript, and T.K. wrote the final manuscript.

## Competing Interest

The authors declare no competing interests.

## Materials & Correspondence

Correspondence and requests for materials should be addressed to T.K.

---

**Extended Data Fig 1.**
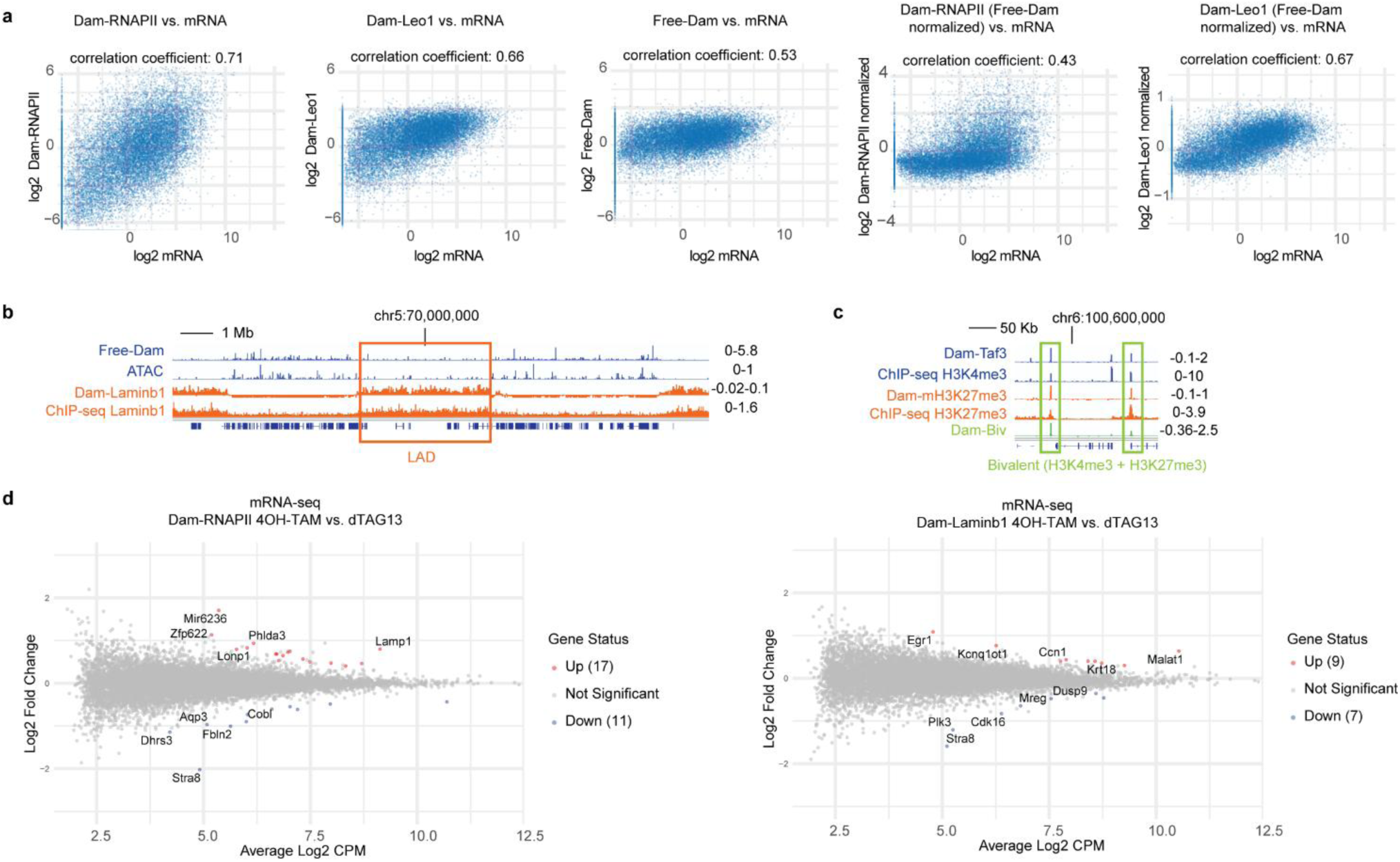
**a**, Scatter plots comparing mRNA-seq against Dam-RNAPII, Dam-Leo1, Free-Dam, Dam-RNAPII (Free-Dam normalized), Dam-Leo1 (Free-Dam normalized) in mESCs. Free-Dam normalization does not increase the Pearson’s correlation between Dam-POIs and mRNA-seq (*n* = 3). **b**, Genome browser views in mESC. Free-Dam and ATAC-seq, which label open chromatin, show mutually exclusive patterns against Dam-Laminb1 and Laminb1 ChIP-seq, which label heterochromatin lamina-associated domains (LADs) (*n* = 2, biological replicates). **c**, Genome browser views in mESC. Dam-Biv labels H3K4me3 (Dam-Taf3 and H3K4me3 ChIP-seq) and H3K27me3 (Dam-mH3K27me3 and H3K27me3 ChIP-seq) double-positive regions (*n* = 2, biological replicates). **d**, MA plot comparing transcriptome of 4OH-TAM (Dam-on) and dTAG-13 (Dam-off) for Dam-RNAPII (left) and Dam-Laminb1 (right). The numbers of differentially expressed genes are indicated (likelihood-ratio test, FDR < 0.01, no fold-change threshold) (*n* = 2, biological replicates).

**Extended Data Fig 2.**
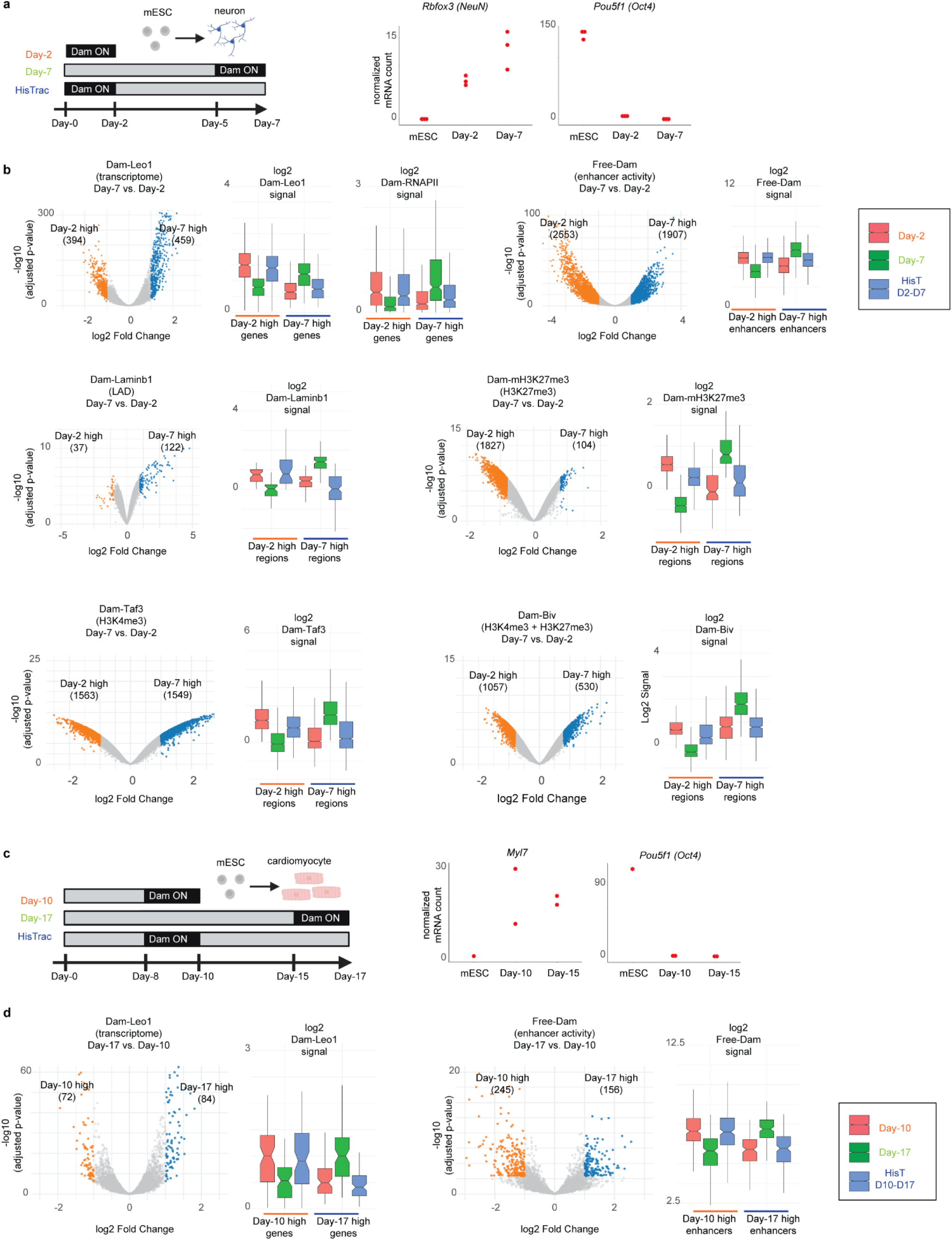
**a**, (left) Timeline of mESC differentiation to neurons with Dam-POI activation windows for Day-2, Day-7, and retrospective HisTrac (Day-2 -> Day-7) labelling. (right) mRNA-seq RPKM (reads per kilobase of exon per million mapped reads) of *Rbfox3* (*NeuN*) and *Pou5f1* (*Oct4*) during neurodifferentiation (*n* = 3). **b**, HisTrac-seq recovers stage-specific m6A signatures for individual Dam-POI during neurodifferentiation (*n* = 2). (left, volcano plot): Differential m6Aenrichment between Day-2 and Day-7 neurodifferentiation snapshots. Each point represents a genomic feature. Genes (Dam-Leo1), enhancers (Free-Dam), 100 Kb bin (Dam-Laminb1), or 10 Kb bin (Dam-mH3K27me3, Dam-Taf3, Dam-Biv) are used. The x-axis shows log2 (fold-change) of Day-2 vs. Day-7 signal; the y-axis shows –log10 (*P*-value) from a likelihood-ratio test. Features significantly enriched at Day-2 are highlighted in orange (“Day-2 high”), and those enriched at Day-7 in blue (“Day-7 high”) (log2FC > 1, FDR < 0.01). (right, boxplots): m6A signal distributions for the Day-2-high and Day-7-high feature sets across three conditions: Day-2 snapshot, Day-7 snapshot, and retrospective HisTrac. For each feature set, boxplots show the median, interquartile range and whiskers to 1.5 × IQR, with individual replicate values overlaid. HisTrac signal closely matches the Day-2 snapshot, demonstrating faithful retrospective recovery of past transcriptional and epigenetic profiles. **c**, (left) Timeline of mESC differentiation to cardiomyocytes. (right) mRNA-seq RPKM of *Mhl7* and *Pou5f1* (*Oct4*) during cardiomyocyte differentiation (*n* = 2). **d**, HisTrac-seq recovers stage-specific m6A signatures for individual Dam-POI in cardiomyocyte differentiation (*n* = 2). Same analysis with **b**.

**Extended Data Fig 3.**
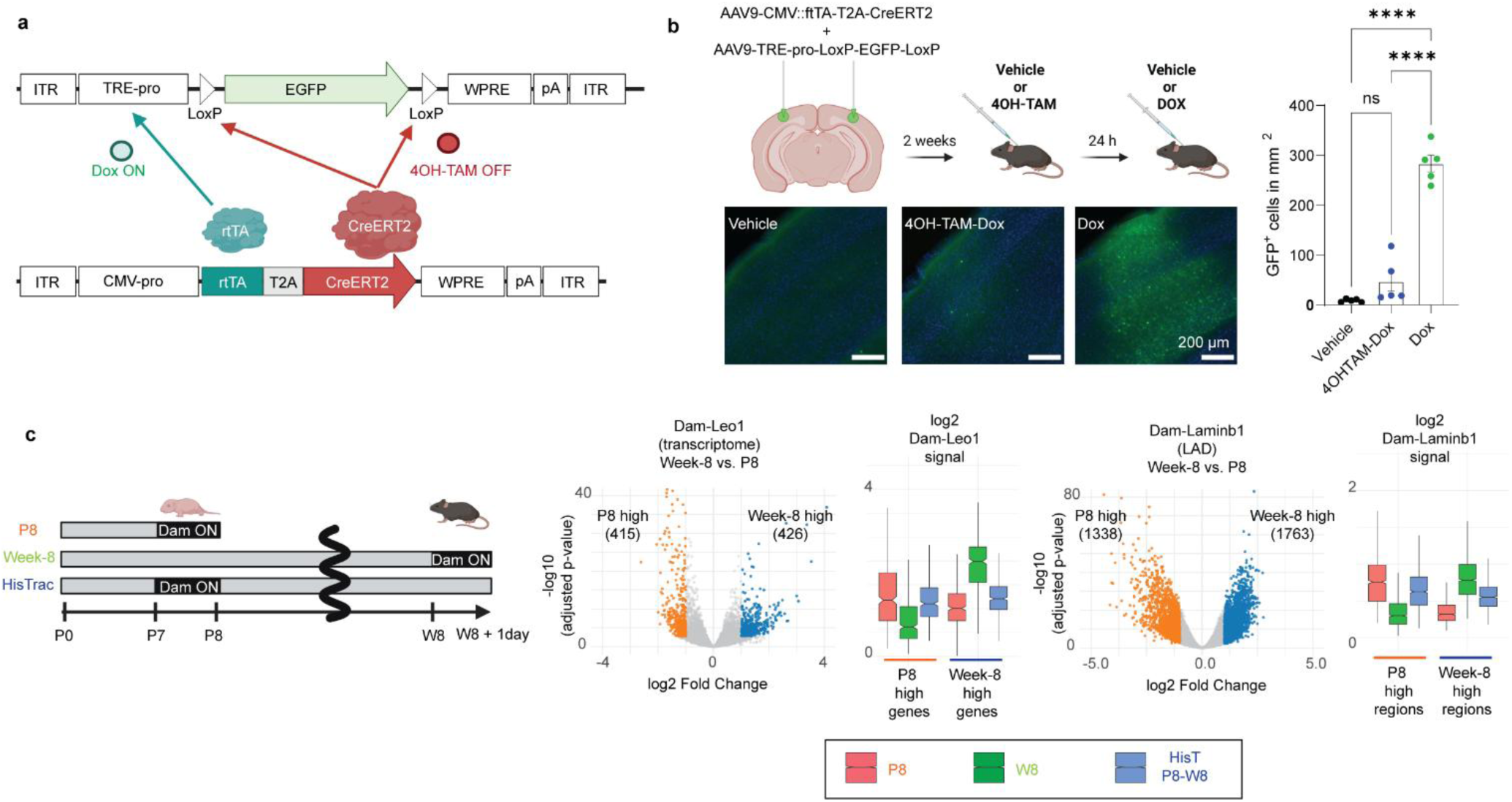
**a**, Schematic of AAV9-mediated delivery of the *TRE-Tight2–LoxP-EGFP–LoxP* reporter and *CMV-rtTA-T2A-CreERT2*. Dox induces EGFP expression via rtTA, whereas 4OH-TAM triggers Cre-LoxP excision of the cassette to prevent expression. **b**, (top left) Following cortical AAV injection, mice were exposed to 4OH-TAM and Dox treatment, and EGFP signal was quantified on slices. (right) Bar plots quantifying EGFP signals from three experimental groups: vehicle only (*n* = 5 different animals), Dox only (*n* = 5 different animals), and 4OH-TAM pretreatment followed by Dox (*n* = 5 different animals). Ordinary One-way ANOVA, F=1,363 (2, 12), *P*-value <0.0001). (bottom left) Representative images of cortical sections. EGFP fluorescence is shown in green; DAPI nuclear stain in blue. **c**, *In vivo* history-tracing workflow and genome-wide m6A profiling (*n* = 2–4). (top) Activation windows for neonatal (P8), adult (Week-8), and retrospective HisTrac (P8 -> week 8) labelling with Dam-Leo1 and Dam-Laminb1. (bottom) Volcano plot: Differential m6A enrichment between P8 and Week-8 snapshots. Each point represents a genomic feature. Genes (Dam-Leo1) or 100 Kb bin (Dam-Laminb1) are used. The x-axis shows log2(fold-change) of P8 vs. Week-8 signal; the y-axis shows –log10 (*P*-value) from a likelihood-ratio test. Features significantly enriched at P8 are highlighted in orange (“P8 high”), and those enriched at Week-8 in blue (“Week-8 high”) (log2FC > 1, FDR < 0.01). Boxplots: m6A signal distributions for the P8 high and Week8 high feature sets across three conditions: P8 snapshot, Week-8 snapshot, and retrospective HisTrac. For each feature set, boxplots show the median, interquartile range, and whiskers to 1.5 × IQR, with individual replicate values overlaid. HisTrac signals closely match the P8 snapshot, demonstrating faithful retrospective recovery of past molecular profiles.

**Extended Data Fig 4.**
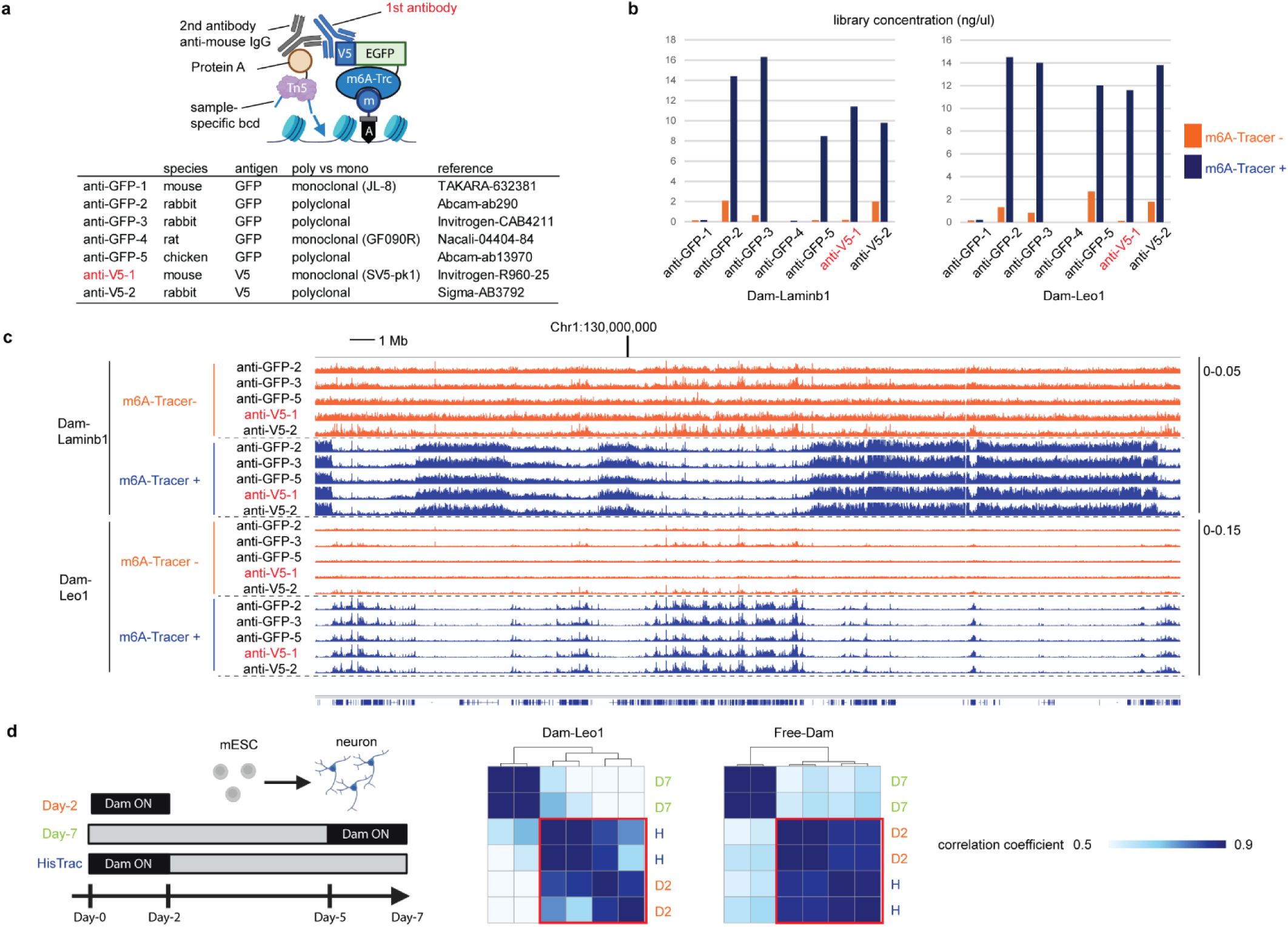
**a**, Screening of primary antibodies against EGFP or V5 epitope tag for Dam&Tag method. Seven antibodies are listed. **b**, Bulk Dam&Tag library yields from Dam-Leo1 or Dam-Laminb1 mESCs with (blue) and without (orange) V5-EGFP-m6A-Tracer (*n* = 1). 300 K cells were used for the starting material, and library concentration was quantified after 10 PCR cycles. Five antibodies showed an increased yield in the presence of a tracer. **c**, Libraries derived from five antibodies were sequenced, and genome browser tracks of m6A signals are visualized. Comparisons between m6A-Tracer-negative (orange) and tracer-positive (blue) conditions revealed that anti-V5 antibody-1 provides the best signal-to-noise ratio for POI-specific signals. Dam-Laminb1 marks heterochromatin in regions mutually exclusive with Dam-Leo1, without Free-Dam normalization. **d**, *In vitro* neurodifferentiation history tracing with Dam&Tag protocol (*n* = 2). (left) Timeline of mESC differentiation to neurons with Dam-POI activation windows. (right) Genome-wide m6A quantification in fixed genomic 10kb bins clustered by Pearson’s correlation.

**Extended Data Fig 5.**
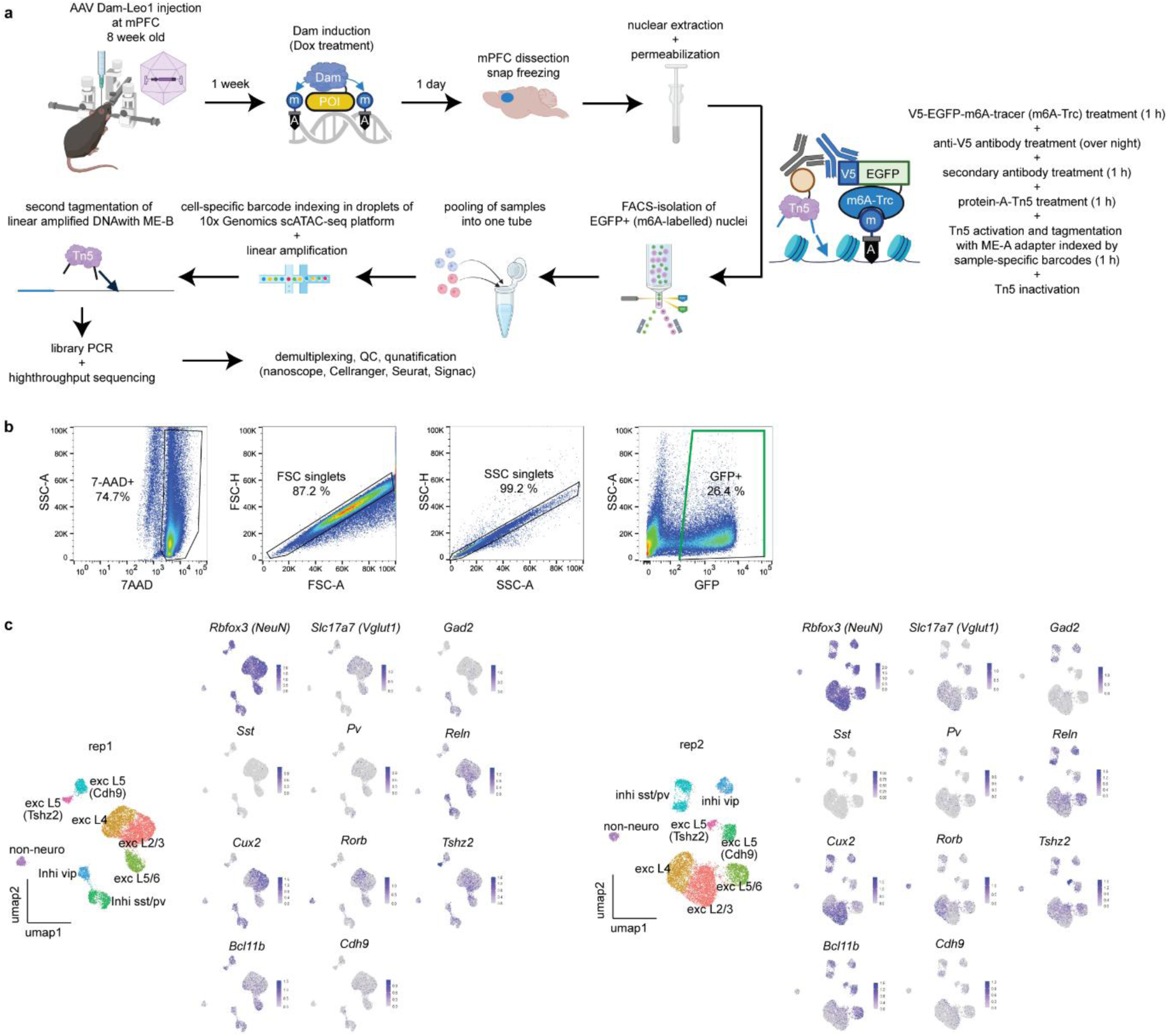
**a**, Schematic of the droplet-based single-cell (sc) Dam&Tag protocol. One day after Dam-POI activation in the mouse cortex, the infected brain region is dissected, and nuclei are extracted. Permeabilized nuclei are incubated with V5-EGFP-m6A-Tracer, mouse anti-V5 antibody, and secondary anti-mouse IgG antibody. Subsequently, nuclei are tagmented by Protein A–Tn5 loaded with ME-A adapters bearing sample-specific barcodes. Following FACS-isolation of nuclei stained with V5-EGFP-m6A-Trace, nuclei are pooled and subjected to 10x Chromium scATAC-seq chemistry. Overloaded nuclei (i.e., multiple per droplet) receive cell-specific barcodes during the standard barcoding reaction. Dual indexing preserves both sample origin and single-cell identity for high-throughput m6A profiling in one Chromium channel. Following linear amplification, secondary tagmentation is carried out with the ME-B adapter. Library preparation PCR, and high-throughput sequencing are carried out. We use publicly available informatics tools for the downstream analysis. **b**, Representative FACS plots from adult mouse cortex expressing Dam-Leo1. 7-AAD signal is used to gate intact nuclei and single events are gated. Nuclei stained with V5-EGFP-m6A-Tracer show an EGFP+ population, which is gated to enrich m6A-marked neurons for downstream scDam&Tag library preparation. **c**, UMAP embedding per replicate with cell type-specific genes.

**Extended Data Fig 6.**
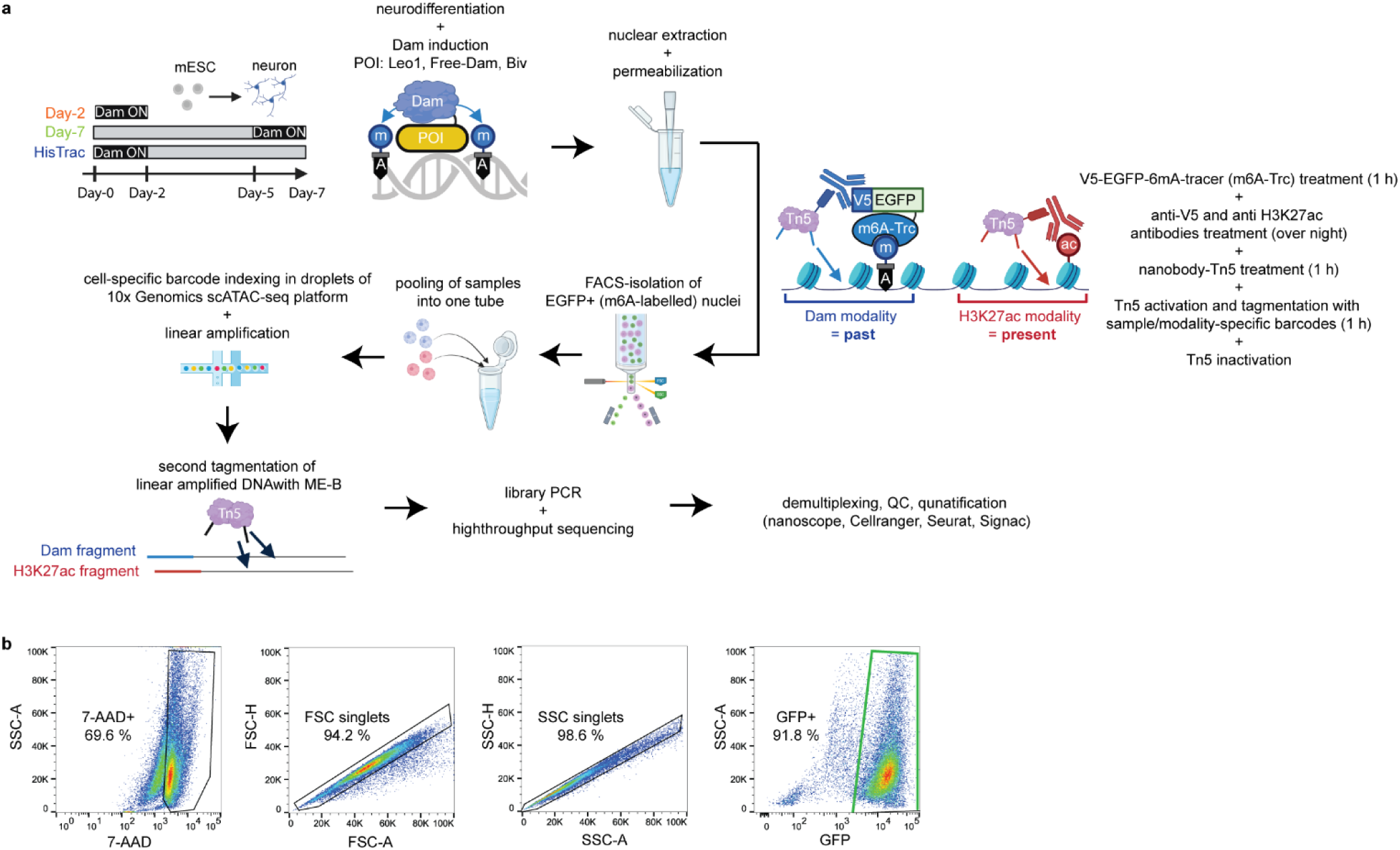
**a**, Schematic of single-cell (sc) HisTrac-seq for simultaneous profiling of past and present molecular profiles. After Dam-POI activation, nuclei are extracted and permeabilized. Permeabilized nuclei are first treated with V5-EGFP-m6A-Tracer to bind m6A marks. Mouse anti-V5 antibody and anti-mouse-IgG nanobody–Tn5 tagment chromatin at historical (m6A) marks. In parallel, rabbit anti-H3K27ac antibody and anti-rabbit-IgG nanobody–Tn5 target active histone modifications to profile present transcriptional activity. ME-A adapters carry modality- and sample-specific barcodes, enabling post-sequencing demultiplexing of m6A versus H3K27ac modalities and sample information. Following FACS-isolation of nuclei stained with V5-EGFP-m6A-Tracer, nuclei are pooled and subjected to 10x Chromium scATAC-seq chemistry. Overloaded nuclei (i.e., multiple per droplet) receive cell-specific barcodes during the standard barcoding reaction. Dual indexing preserves sample origin, modality, and single-cell identity for high-throughput m6A profiling in one Chromium channel. Following linear amplification, secondary tagmentation is carried out with the ME-B adapter. Library preparation PCR and high-throughput sequencing are carried out. We use publicly available informatics tools for the downstream analysis. **b**, Representative FACS plots from mESC-derived neurons expressing Dam-Leo1. Nuclei stained with V5-EGFP-m6A-Tracer show an EGFP+ population, which is gated to enrich m6A-marked neurons for downstream library preparation. 7-AAD signal is used to gate intact nuclei.

**Extended Data Fig. 7.**
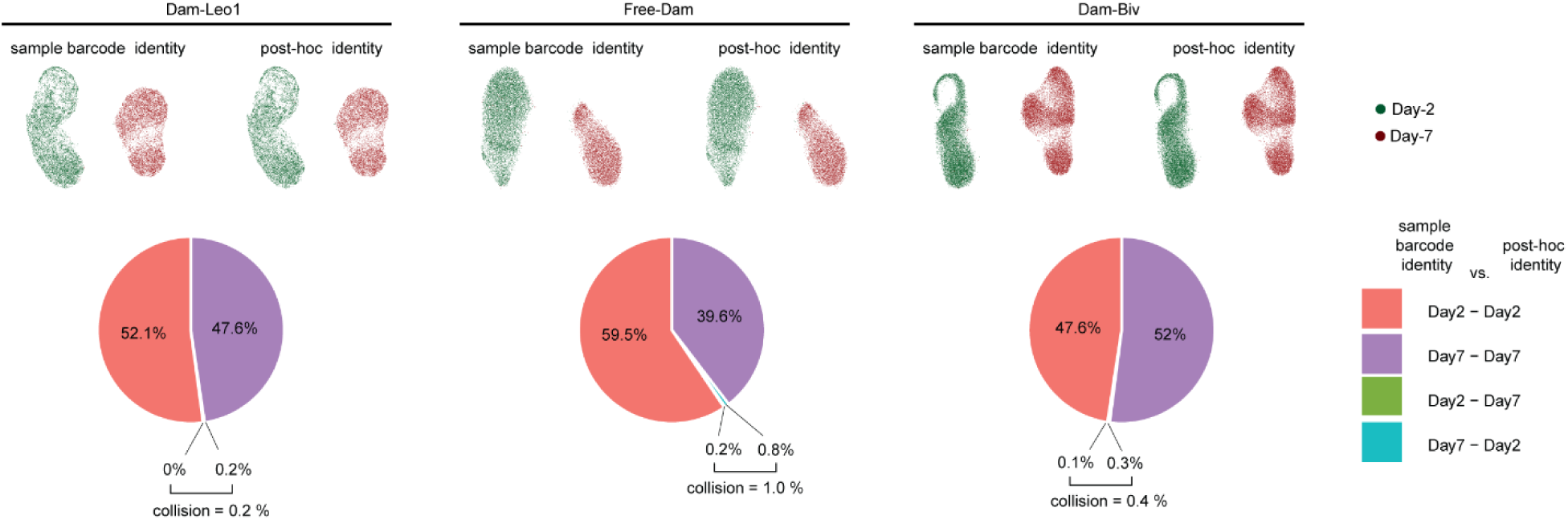
Evaluation of Tn5 barcode hopping in scDam&Tag. (top) UMAP embeddings of H3K27ac profiles for Day-2 (green) and Day-7 (red) cells, shown separately for Dam-Leo1 (left), Free-Dam (middle), and Dam-Biv (right). Cell colors reflect their sample-barcode assignments and post-hoc clustering identities. (bottom) Pie charts indicating the percentage of cells whose post-hoc clustering-based identity conflict with their sample-barcode labels. Collision rates are 0.2 % for Dam-Leo1, 1.0 % for Free-Dam, and 0.4 % for Dam-Biv, demonstrating negligible Tn5 barcode hopping.

**Extended Data Fig. 8.**
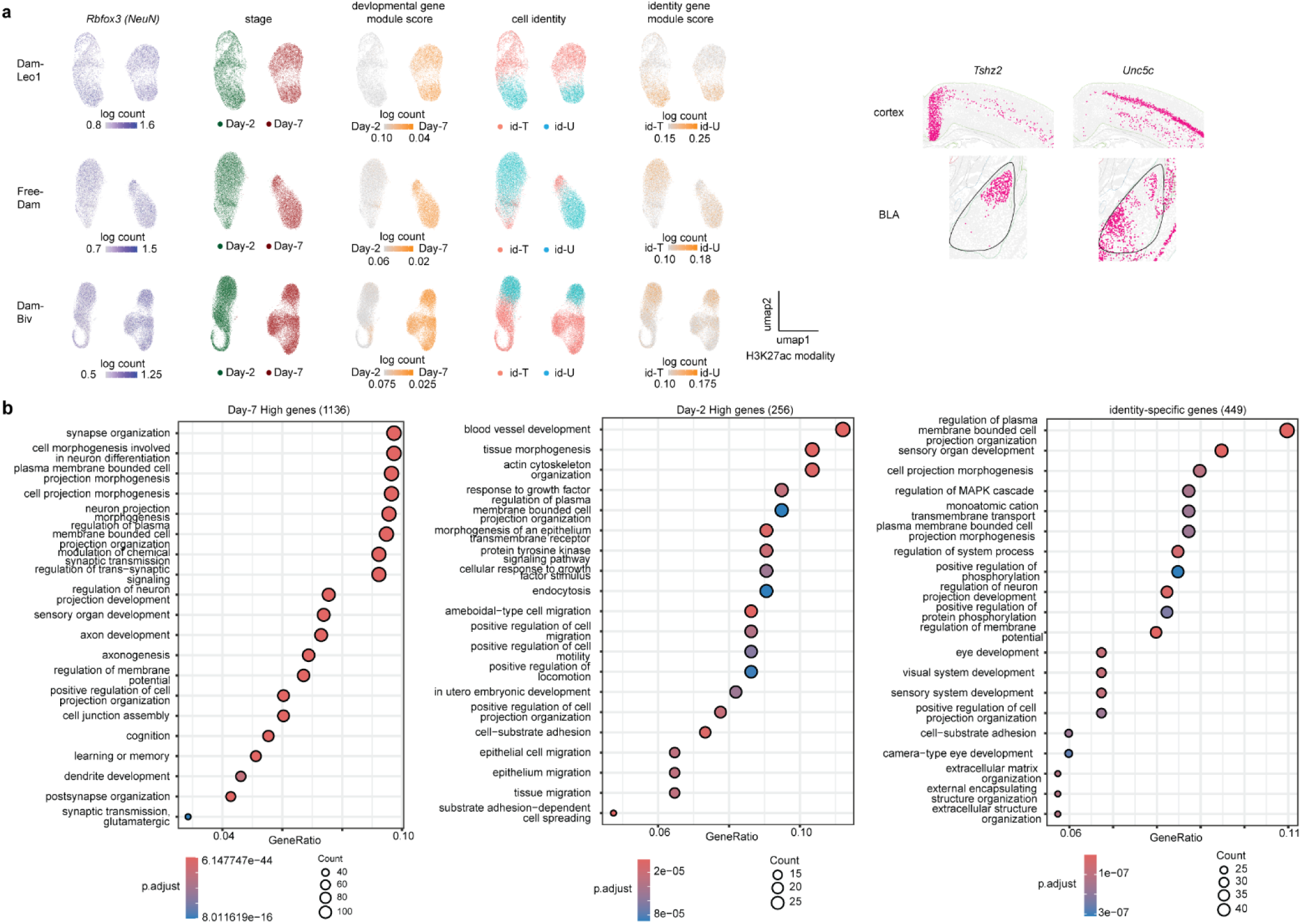
**a**, (left) UMAP embedding of H3K27ac modality shown separately for Dam-Leo1 (top), Free-Dam (middle), and Dam-Biv (bottom). Cells primarily cluster by developmental stages, namely Day-2 and Day-7. They are further subdivided into two identities, namely id-Tshz2 (id-T) and id-Unc5c (id-U). They are all *Rbfox3* (*NeuN*)-positive postmitotic neurons. Developmental and identity gene module scores are also shown. The sample size is indicated in Fig. 4c. (right) Coronal section of adult mouse brain imaged in the Allen Brain Cell Atlas. Red dots indicate excitatory neurons expressing *Tshz2* and *Unc-5c* in the cortex and basolateral amygdala. While *Tshz2* is expressed in the medial cortex and lateral amygdala, *Unc5c* is expressed in the lateral cortex and basal amygdala. **b**, Gene ontology analysis of Day7-high genes, Day-2 high genes, and identity-specific genes, identified by the analysis of single-cell data. The sample size is indicated in Fig. 4c.

**Extended Data Fig. 9.**
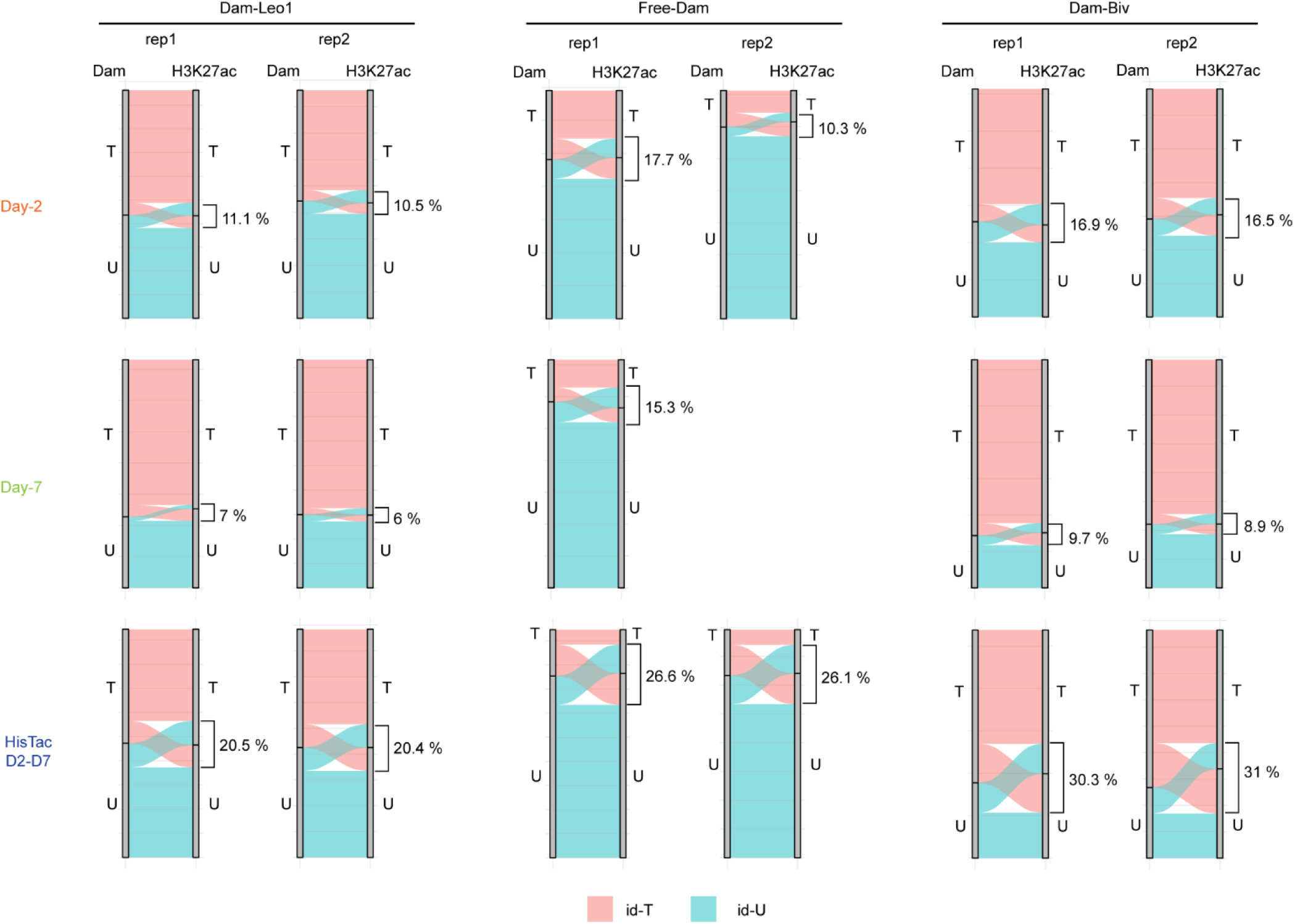
Alluvial plots quantify cell identity transition (i.e., id-T vs. id-U) between Dam and H3K27ac modalities for individual replicates (see Fig. 5b for the replicate-merged version). The discrepancies between the two modalities are highlighted. Discrepancy rates for HisTrac-seq far exceed the background discrepancy rates for Day-2 and Day-7 snapshots, indicating bona fide disruptive “identity jumps” during differentiation.

**Extended Data Fig. 10.**
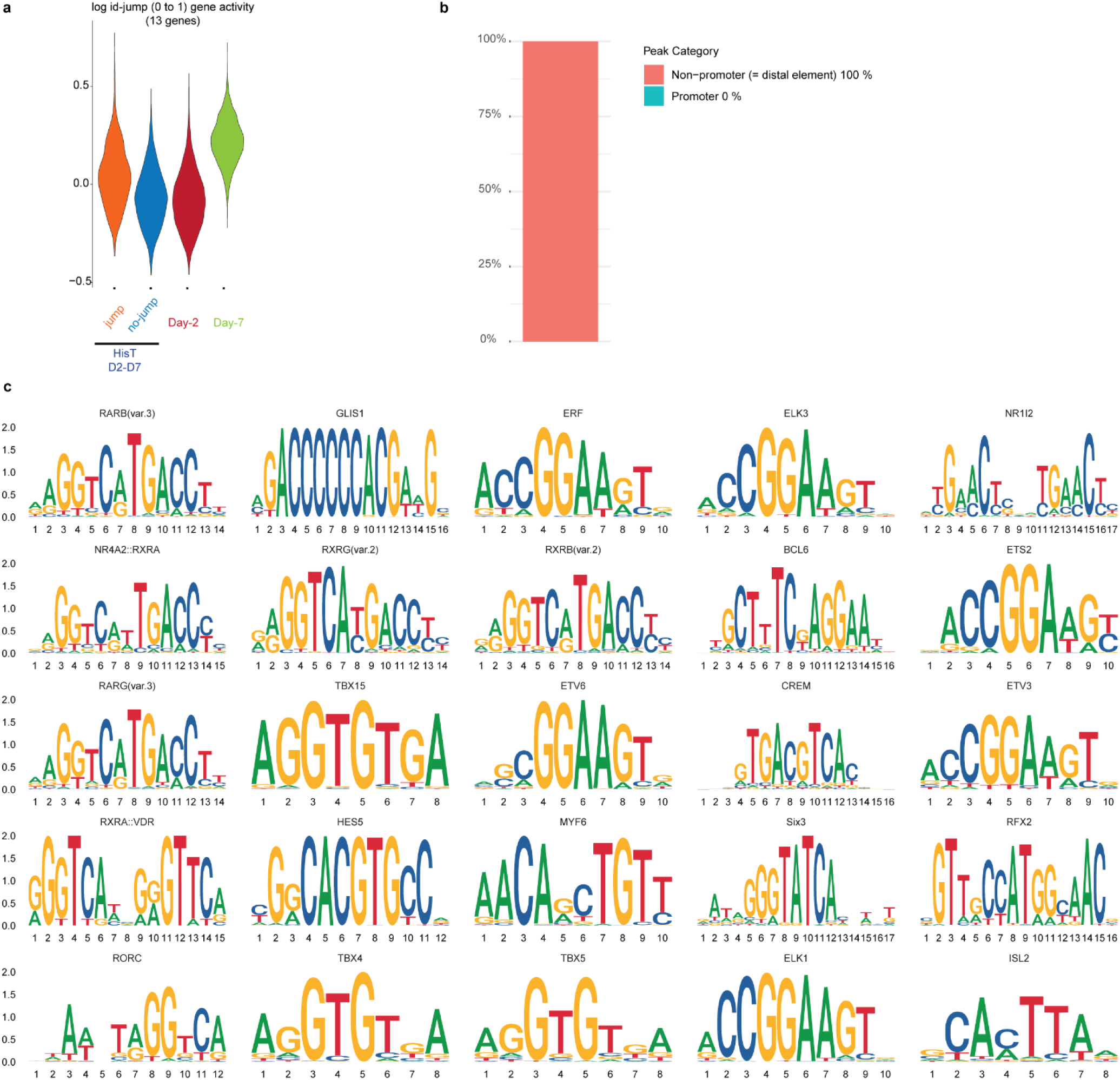
**a**, Violin plots of Dam-Leo1-measured expression for the 13 genes associated with id-T to id-U identity jumps. Cells that undergo jumps (orange) versus those that maintain identity (blue) are compared. These genes normally show higher expression at Day-7 (green) versus Day-2 (dark red) (*n* = 2). **b**, Ratio comparing promoter vs. distal element for the 141 peaks associated with identity jumps (Fig. 6c). **c**, Transcription factor–motif enrichment analysis in the 141 jump-associated enhancers, with the top 25 most significantly enriched (*n* = 2).

**Extended Data Fig. 11.**
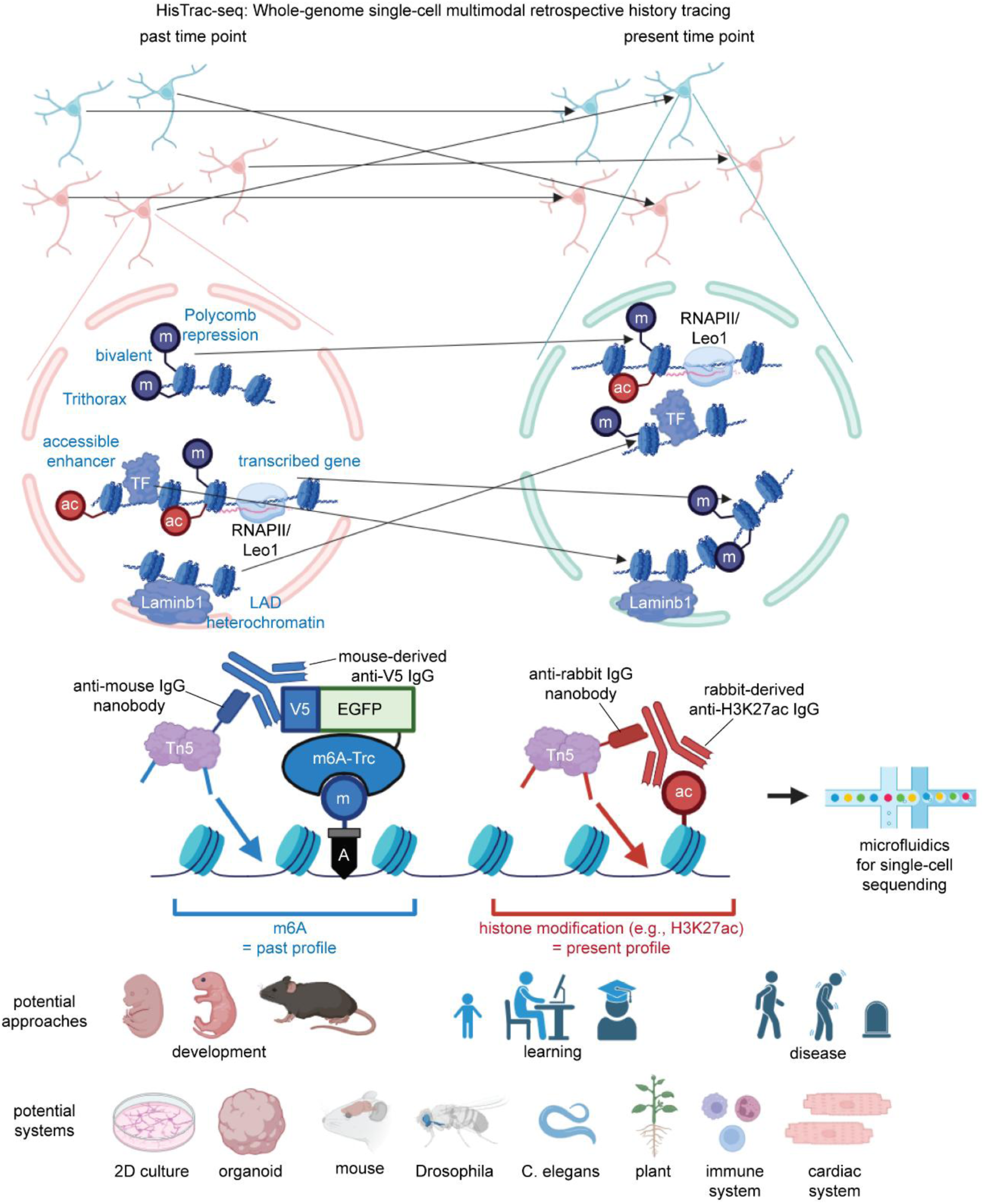
Summary diagram. In contrast to conventional snapshot sequencing, HisTrac-seq simultaneously maps the past and present whole-genome profiles at the single-cell resolution. This allows us to trace how individual cells develop their molecular profiles over time. This can interrogate multiple regulatory layers, including transcriptome, chromatin accessibility, heterochromatin repression, Polycomb repression, promoter activity, and bivalent poised state. HisTrac-seq is a “temporal multi-omics” platform potentially applicable to diverse biological questions and experimental systems.

## Notes

### Competing Interest Statement

The authors have declared no competing interest.

